# Longitudinal White Matter Trajectories in Clinical High Risk and First-Episode Psychosis: Findings from the Multi-Centre PSYSCAN Study

**DOI:** 10.64898/2026.07.20.739516

**Authors:** Lara Bolte, George Gifford, Mauricio Serpa, Michiel Cottaar, Paola Dazzan, Paolo Fusar-Poli, Stefania Tognin, Matthew Kempton, Inge Winter-von Rossum, Margot I. E. Slot, Hendrika van Hell, Arija Maat, Lieuwe de Haan, Benedicto Crespo Facorro, Birte Glenthøj, Stephen Lawrie, Colm McDonald, Thérèse van Amelsvoort, Celso Arango, Irina Falkenberg, Barnaby Nelson, Silvana Galderisi, Rodrigo A. Bressan, Jun Soo Kwon, Kang Ik Kevin Cho, Mark Weiser, Romina Mizrahi, Gabriele Sachs, Matthias Kirschner, Maxime Taquet, Dominic Oliver, PSYSCAN Consortium, Rene Kahn, Philip McGuire

## Abstract

Psychosis has been linked to changes in diffusion-derived fractional anisotropy (FA) across multiple white matter tracts, yet most evidence is cross-sectional. Longitudinal findings are inconsistent, and the clinical relevance of white matter changes over time remains unclear. To address this gap, we investigated longitudinal white matter changes and their clinical correlates in early psychosis in a large, multi-centre diffusion-tensor imaging study.

Across 18 sites, 407 participants (healthy controls: *n* = 97, 59.8% men, mean age 23.9±4.6 years; clinical high risk of psychosis: *n* = 158, 53.5% men, mean age 23.0±4.9 years; first-episode psychosis: *n* = 152, 71.1% men, mean age 25.1±5.4 years) were scanned at up to three time points over 12 months. FA was assessed in the cingulum bundle, the superior longitudinal fasciculus, the inferior fronto-occipital fasciculus, and at the whole brain level. FA trajectories were analysed using linear mixed-effects models to test effects of group, social and occupational functioning, and (attenuated) psychotic symptoms, while controlling for demographic and socioeconomic covariates.

No significant group differences in FA were observed, either globally or within tracts (*p* > .05). Longitudinal FA trajectories were not associated with changes in (attenuated) psychotic symptoms or functioning (*p* > .05). In contrast, higher baseline antipsychotic medication dose was significantly associated with lower FA (*p_corr_* < .05).

Overall, these findings indicate that white matter microstructure is relatively stable during the clinical-high-risk and first-episode phases of psychosis, and that it is not clearly associated with variation in symptom severity or functional outcomes over time.

## Introduction

Converging evidence from genetic, post-mortem, and neuroimaging studies suggests that disruptions in white matter (WM) represent a neurobiological feature of psychotic disorders. Genetic studies show that the polygenic risk score for schizophrenia is associated with WM alterations, with some risk variants linked with biological processes relevant to myelination ^1–3^. Consistent with this, post-mortem findings report reduced oligodendrocyte density in chronic schizophrenia ^4^. Neuroimaging studies in individuals with psychosis further demonstrate widespread WM disruptions, including reduced neurite density and increased extracellular free water in first-episode psychosis (FEP) and chronic schizophrenia ^5–8^.

Among neuroimaging approaches, diffusion tensor imaging (DTI) has provided some of the strongest evidence for WM alterations in psychotic disorders and their prodromal stage, with abnormalities reported in mean, radial, and axial diffusivity ^9–11^. The most replicable and robust DTI-derived metric is fractional anisotropy (FA), an indirect proxy of WM microstructure that quantifies the directional movement of water molecules, ranging from 0 (isotropic diffusion) to 1 (diffusion restricted to a single orientation).

Cross-sectional DTI studies consistently suggest that individuals at clinical high-risk for psychosis (CHR-P) and patients with FEP exhibit widespread reductions in FA relative to healthy controls (HC), with more pronounced reductions in FEP than CHR-P ^10,12–16^. The most consistently implicated pathways in CHR-P and FEP are long-range association tracts connecting frontal regions with parietal, temporal, and limbic cortices, including (1) the cingulum bundle (CB), (2) the superior longitudinal fasciculus (SLF), and (3) the inferior fronto-occipital fasciculus (IFOF). These tracts are thought to support cognitive processes that are disrupted in psychosis, such as executive functioning, language, memory and self-monitoring, underscoring their clinical relevance ^17–19^. Altered WM microstructure has also been linked to clinical outcomes in the early stages of psychosis. In FEP, FA has been linked to treatment response ^20^, while in CHR-P, WM alterations have been associated with subsequent transition to psychosis ^21^. Associations between FA and positive and negative symptom severity have been reported in both CHR-P and FEP ^22–24^, although findings remain heterogeneous across studies ^13^. Notably, relevant tracts again include the CB, SLF, and IFOF ^20,24^. Together, these findings suggest that disruptions to WM microstructure in the early stages of psychosis, particularly in these pathways, may contribute not only to neurobiological vulnerability but also to clinically meaningful variation in illness expression.

Despite this, the trajectory of WM changes over time in CHR-P and FEP remains poorly understood, as relatively few studies have examined longitudinal changes in WM microstructure. A recent systematic review ^14^ reported that FA may increase over time in CHR-P groups, in some cases approaching levels observed in HC ^25^. In contrast, another review ^12^ described largely stable or declining FA in individuals with CHR-P ^26^. These inconsistent findings may partly reflect limitations of the existing literature, including small sample sizes and incomplete control of potentially confounding factors such as antipsychotic exposure. Larger and more rigorously controlled longitudinal studies are therefore needed to clarify the trajectory of WM microstructure in the prodromal and early stages of psychosis.

The present study sought to investigate longitudinal changes in WM microstructure in CHR-P and FEP compared with HC using a prospective, multi-centre study design. Participants underwent DTI at up to three time points over 12 months. Based on prior findings in psychosis, we focused our analyses on the CB, SLF, and IFOF, and global FA. We first tested the hypothesis that FA remains largely stable in HC, with modest reductions in FA seen in CHR-P that are more pronounced in FEP. Our second hypothesis was that, within the CHR-P and FEP groups, greater reductions in FA would be associated with greater severity of psychotic symptoms and poorer levels of functioning. We expected these associations to be evident both globally and within the CB, SLF, and IFOF. Analyses were not designed to examine transition to psychosis within the CHR-P group, as the current analytic framework is underpowered for such subgroup comparisons.

## 1. Methods

### 1.1. Study Design

Participants were recruited to an international, longitudinal, multi-centre prospective cohort study (PSYSCAN) between July 2016 and December 2019 across 18 sites: Australia (Melbourne), Austria (Vienna), Denmark (Copenhagen), Germany (Marburg), Ireland (Galway), Israel (Ramat Gan), Italy (Naples), The Netherlands (Amsterdam, Maastricht, Utrecht), Spain (Santander, Madrid), Switzerland (Zurich), United Kingdom (Edinburgh, London), Canada (Toronto), South Korea (Seoul), and Brazil (Sao Paulo). Detailed descriptions of the study design and cohorts have been published elsewhere ^27–29^.

### 1.2. Participants

As part of the PSYSCAN study, participants included HC, CHR-P, and FEP (Table 1). CHR-P status was defined by the Comprehensive Assessment of At-Risk Mental States (CAARMS) ^31^, or the ‘basic symptoms’ criteria using the Schizophrenia Proneness Instrument (SPI-A) ^32^. FEP status was defined by a DSM-IV diagnosis of schizophrenia, schizoaffective disorder, or schizophreniform disorder, with psychosis onset (first contact with a healthcare professional at which a diagnosis of a psychotic disorder was made) within three years of study entry.

**Table 1.** Sociodemographic Variables per Group.

|  | HC | CHR | FEP |
| --- | --- | --- | --- |
| Total |  |  |  |
| *N* | 97 | 158 | 152 |
| Age in years |  |  |  |
| Mean (*SD*) | 23.9 (4.6) | 23.0 (4.9) | 25.1 (5.4) |
| Gender |  |  |  |
| Male *n* (%) | 58 (59.8 %) | 84 (53.5 %) | 108 (71.1 %) |
| Ethnicity |  |  |  |
| African | 7 | 16 | 28 |
| Asian Indian | 5 | 4 | 4 |
| East Asian | 20 | 29 | 1 |
| Inter Racial | 1 | 3 | - |
| Middle Eastern | 1 | 3 | 2 |
| Other | 3 | 9 | 10 |
| White | 60 | 93 | 107 |
| WAIS |  |  |  |
| Mean (*SD*) | 113.3 (14.7) | 106.4 (17.4) | 91.9 (19.2) |
| Antipsychotic dosage mg/day |  |  |  |
| Mean (*SD*) | 0 (0) | 15.8 (71.5) | 224.9 (201.2) |

*Note.* One participant reported a non-binary gender; to reduce identifiability due to small cell size, this value was coded as missing. Ethnicity data were missing for one participant, and antipsychotic dosage (chlorpromazine equivalents) was missing for 13 participants. No other data were missing. HC = Healthy Controls; CHR = Clinical High-Risk; FEP = First-Episode Psychosis.

Exclusion criteria for any participant group were: (i) previous neurosurgery or neurological disorder, (ii) IQ < 70, (iii) incapability to comprehend the purpose of the study or to make an informed decision on study participation, (iv) a history of a head injury resulting in unconsciousness for > 1 hour, (v) ages outside of 16 – 40 years (except for one site, Madrid, which recruited 14 – 40), and (vi) any contraindications for undergoing magnetic resonance imaging (MRI), including pregnancy. Specific to HC, exclusion criteria further included (i) a lifetime history of any DSM-IV Axis-I or Axis-II mental disorder (borderline, paranoid and schizotypal personality disorder), (ii) a first-degree relative with affective or non-affective psychosis, (iii) meeting CHR-P criteria, and (iv) a lifetime history of antipsychotic or psychoactive medication. Additional exclusion criteria for the CHR-P group were (i) more than 30 days cumulative exposure to antipsychotic medication at a dosage for treating FEP in the 3 months prior to baseline, and (ii) any past episode of frank psychosis lasting more than 7 days.

Participants, or their legal representatives for participants < 18 years old, gave written informed consent. Ethical approval was granted by local research ethics committees. All procedures complied with ethical standards of the relevant national and institutional committees on human experimentation and with the Helsinki Declaration of 1975 (revised in 2008).

### 1.3. Clinical Assessments

Participants completed clinical assessments at all scanning time points: baseline, 6-month, and 12-month follow-up, apart from FEP, who did not have a 6-month follow-up scanning time point. Functioning was assessed in all participants using the Social and Occupational Functioning Assessment Scale (SOFAS) ^33^. Psychotic symptom severity was assessed in CHR-P using the CAARMS ^31^ and in FEP with the Positive and Negative Syndrome Scale (PANSS) ^34^ (Table 2). For both clinical measures, total positive symptom severity sum scores were used.

**Table 2.**
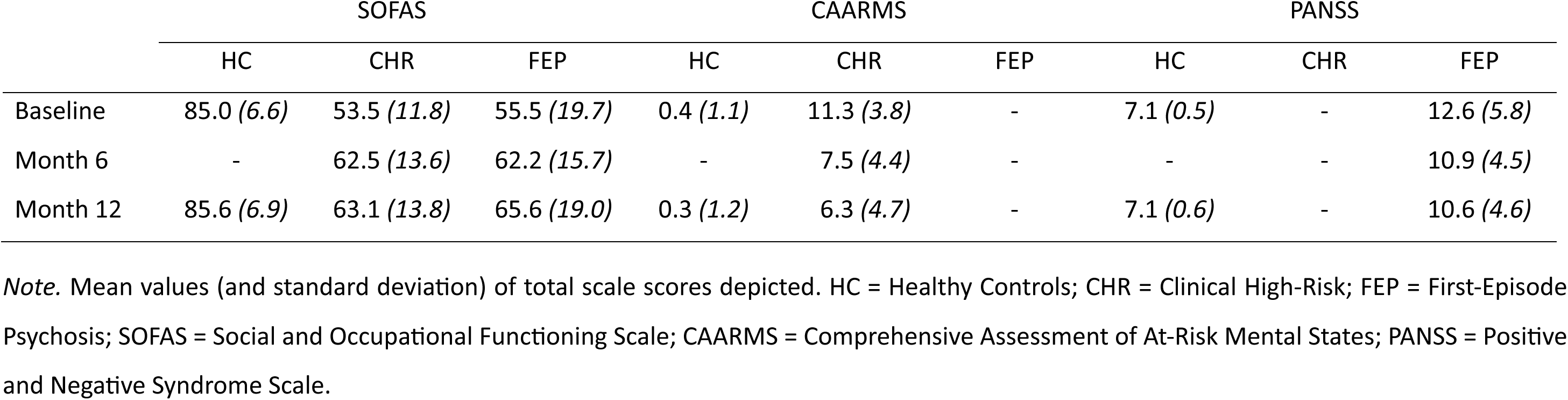
Clinical Characteristics per Group per Timepoint.

|  | SOFAS |  |  | CAARMS |  |  | PANSS |  |  |
| --- | --- | --- | --- | --- | --- | --- | --- | --- | --- |
|  | HC | CHR | FEP | HC | CHR | FEP | HC | CHR | FEP |
| Baseline | 85.0 (6.6) | 53.5 (11.8) | 55.5 (19.7) | 0.4 (1.1) | 11.3 (3.8) | - | 7.1 (0.5) | - | 12.6 (5.8) |
| Month 6 | - | 62.5 (13.6) | 62.2 (15.7) | - | 7.5 (4.4) | - | - | - | 10.9 (4.5) |
| Month 12 | 85.6 (6.9) | 63.1 (13.8) | 65.6 (19.0) | 0.3 (1.2) | 6.3 (4.7) | - | 7.1 (0.6) | - | 10.6 (4.6) |

*Note.* Mean values (and standard deviation) of total scale scores depicted. HC = Healthy Controls; CHR = Clinical High-Risk; FEP = First-Episode Psychosis; SOFAS = Social and Occupational Functioning Scale; CAARMS = Comprehensive Assessment of At-Risk Mental States; PANSS = Positive and Negative Syndrome Scale.

### 1.4. Diffusion Tensor Imaging

#### 1.4.1. Data Acquisition

Diffusion tensor imaging (DTI) data were obtained using 3 T scanners from Siemens, Philips, or GE equipped with 8 – 64 channel head coils (Table S1). All sites used echo-planar imaging covering the whole brain with axial slice orientation, slice thickness was 2.5 mm, reconstructed at an in-plane resolution of 2.5 x 2.5 mm^2^ (except Ramat Gan, 1.25 x 1.25 mm^2^); b value = 1000 s/m^2^; one or five interspersed b = ∼0s/mm^2^ images; 64 non-collinear diffusion gradient directions (except Sao Paulo, 32); 64 slices (except Madrid 62); echo time = 68.3 - 89 ms; no field maps were acquired.

#### 1.4.2. Preprocessing and Quality Control

Brain extraction for all raw DTI scans was performed using FreeSurfer’s SynthStrip tool ^35,36^. Next, a synthetic undistorted b*0* image was generated using Synb0 DisCo ^37,38^ and used together with the distorted b*0* image in TOPUP (FSL) to estimate and correct susceptibility-induced distortions ^39,40^. Motion and eddy-current correction were carried out using FSL’s EDDY ^41^. Quality control was conducted using QUAD and SQUAD, during which we assessed contrast-to-noise ratio, signal-to-noise ratio, and relative motion metrics and visually inspected each scan for inaccuracies ^42^. As a result, 16 scans were excluded.

#### 1.4.3. Fractional Anisotropy Extraction of White Matter Tracts

Following the completion of the preprocessing pipeline, global FA values were acquired from each scan by running DTIFIT ^40^, followed by Tract-Based Spatial Statistics (TBSS) ^43^. Tract-specific regions of interest were obtained by overlaying the global FA skeleton with the John Hopkins University Probabilistic Tractography Atlas ^44^: Cingulum Bundle (CB; mean FA of ‘Cingulum – cingulate gyrus – Left’, ‘Cingulum – cingulate gyrus – Right’, ‘Cingulum – hippocampus – Left’, and ‘Cingulum – hippocampus – Right’), Inferior fronto-occipital fasciculus (IFOF; mean FA of ‘Inferior fronto-occipital fasciculus – Left’ and ‘Inferior fronto-occipital fasciculus – Right’), and Superior longitudinal fasciculus (SLF; mean FA of ‘Superior longitudinal fasciculus – Left’, ‘Superior longitudinal fasciculus – Right’, ‘Superior longitudinal fasciculus – temporal part – Left’, and ‘Superior longitudinal fasciculus – temporal part – Right’; Figure 1).

**Figure 1.**
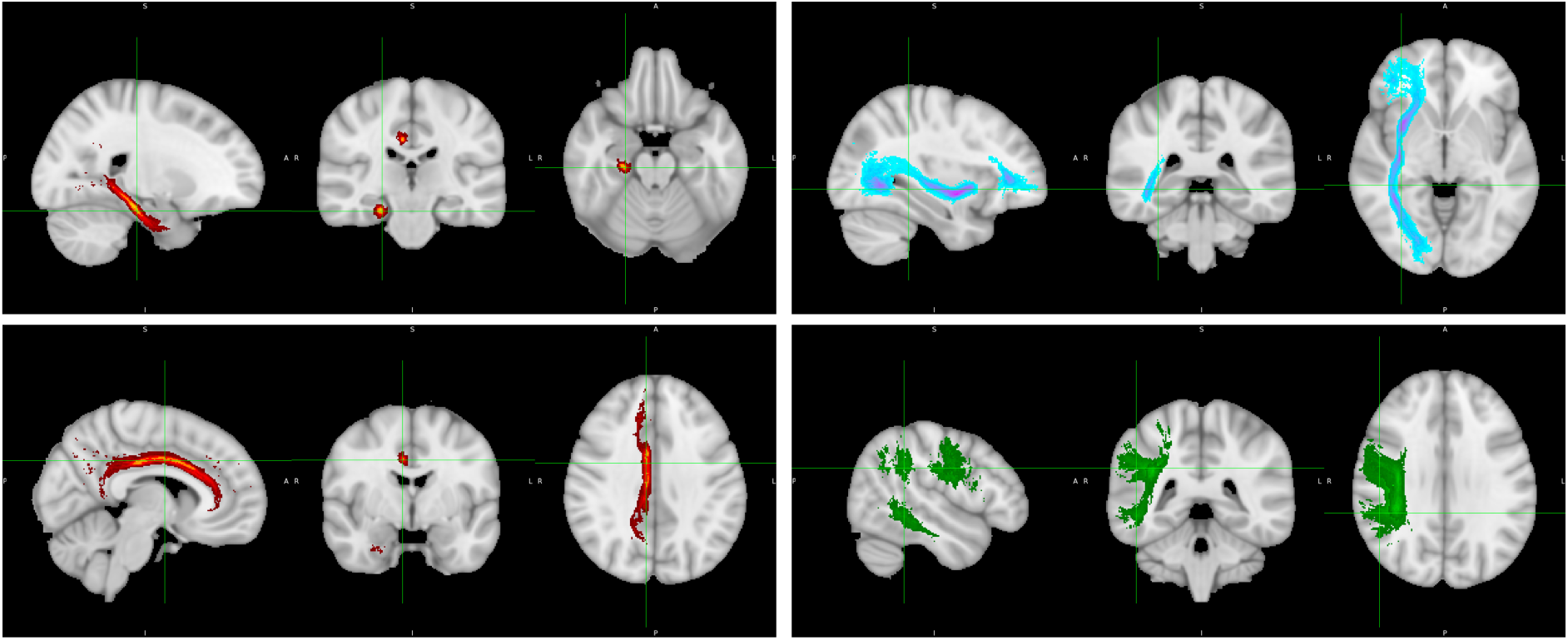
White Matter Tracts. *Note.* Red = right Cingulum Bundle (including the cingulate gyrus and hippocampus), blue = right Inferior Fronto-Occipital Fasciculus, green = right Superior Longitudinal Fasciculus; based on the John Hopkins University Probabilistic Tractography Atlas in FSL.

#### 1.4.4. Site Harmonisation

An additional ‘travelling heads’ (TH) group of healthy volunteers (*n* = 6) was assessed using the study DTI protocol across six sites that contributed the majority of scans (Melbourne, Amsterdam, Maastricht, London, Toronto, and Seoul; 71.3% of all scans). Each TH participant was scanned at all six sites. We implemented mean-offset correction for all scans from these sites using the following formula:

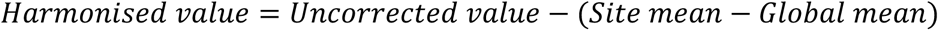

Next, we applied LongCombat ^45^ to the full dataset, controlling for age, ethnicity, gender, group status, transition to psychosis, timepoint, chlorpromazine equivalent antipsychotic dose (capped at 1000 mg/ day), SOFAS score, Scale of Psychosis-risk Syndromes (SOPS) score (derived from CAARMS and PANSS scores; Table S2) ^46,47^ and IQ. TH sites were considered a single mega-site following mean-offset correction.

### 1.5. Statistical Analyses

Statistical analyses were performed in R (version 4.5.2) using the *lme4* package (version 1.1.37). To test our first hypothesis that FA trajectories would differ by group (with stable FA in HC, modest reductions in CHR-P, and more pronounced reductions in FEP), linear mixed-effects models were fitted on the whole dataset to examine the associations between group status (independent variable) and FA (dependent variable) over time at the global and tract-specific levels (CB, SLF, IFOF). We tested for group, time point (baseline, 6-month, and 12-month follow-up), and group-by-time point interaction effects on FA, as fixed effects, while participant-specific random intercepts were included to account for repeated measures. All models were controlled for age, gender, ethnicity, baseline antipsychotic dose (chlorpromazine equivalent), and baseline IQ as fixed effects.

To test our second hypothesis that greater reductions in FA would be associated with a worse clinical profile, analogous linear mixed-effects models were fitted to assess associations between FA trajectories and longitudinal clinical measures, including (i) level of functioning (SOFAS), (ii) severity of psychotic symptoms in FEP (PANSS), and (iii) severity of attenuated psychotic symptoms in CHR-P (CAARMS), with each clinical measure examined in a separate model.

Sensitivity analyses were conducted to evaluate the robustness of the primary group effects. To do so, we (a) divided WM tracts into left and right hemisphere components, (b) stratified the dataset by gender, and (c) excluded baseline antipsychotic dose as a covariate.

Additional post hoc analyses were conducted to facilitate accurate interpretation of the findings, including gender-stratified comparisons of symptom severity and baseline antipsychotic dose, as well as bivariate correlations between symptom severity and baseline antipsychotic dose.

Ninety-five % confidence intervals were obtained via Wald approximation. False discovery rate correction using the Benjamini-Hochberg method was applied to adjust for multiple comparisons across the different regions of interest (global FA, CB, SLF, IFOF). Significance was assessed at a .05 level.

## 2. Results

### 2.1. Participant Characteristics

Longitudinal DTI data were available for a total of *N* = 407 participants included in the present analysis: HC (n = 97, 59.8% men, mean age 23.9±4.6 years); CHR-P (*n* = 158, 53.5% men, mean age 23.0±4.9 years); and FEP (*n* = 152, 71.1% men, mean age 25.1±5.4 years). These data comprised 962 diffusion tensor imaging scans that were acquired at baseline, 6-months, and 12-months follow-up (HC 96/78/80; CHR-P 153/131/119; FEP 152/2/151).

### 2.2. Group Differences in FA Trajectory

There was no evidence of significant differences in FA trajectories between groups, either globally or in specific WM tracts (e.g., global FA [CHR-P vs HC]: β = −0.0021, 95% CI [−0.0069, 0.0028], *p_corr_* = 72; IFOF [FEP vs HC]: β = −0.0027, 95% CI [−0.0095, 0.0040], *p_corr_* = .92). Similarly, neither the main effect of time (e.g., CB [Month 12 vs Baseline]: β = -0.0014, 95% CI [-0.0041, 0.0013], *p_corr_* = .31), nor the group-by-time interaction (e.g., SLF: CHR-P vs HC – Month 6 vs Baseline: β = -0.0022, 95% CI [-0.0054, 0.0010], *p_corr_* = .24) reached statistical significance for any region of interest (Table 3; Figure 2).

**Table 3.** Mixed-Effects Estimates of Group and Time Effects on FA.

| Effect | Global | CB | SLF | IFOF |
| --- | --- | --- | --- | --- |
| Group |  |  |  |  |
| CHR vs HC | -0.0021 ($p_{\text{corr}} = .72$) | -0.0016 ($p_{\text{corr}} = .72$) | -0.0010 ($p_{\text{corr}} = .72$) | -0.0010 ($p_{\text{corr}} = .72$) |
| FEP vs HC | -0.0020 ($p_{\text{corr}} = .92$) | -0.0010 ($p_{\text{corr}} = .92$) | -0.0003 ($p_{\text{corr}} = .92$) | -0.0027 ($p_{\text{corr}} = .92$) |
| Timepoint |  |  |  |  |
| Month 6 | 0.0003 ($p_{\text{corr}} = .90$) | 0.0002 ($p_{\text{corr}} = .90$) | 0.0003 ($p_{\text{corr}} = .90$) | -0.0005 ($p_{\text{corr}} = .90$) |
| Month 12 | -0.0014 ($p_{\text{corr}} = .31$) | -0.0014 ($p_{\text{corr}} = .31$) | -0.0013 ($p_{\text{corr}} = .31$) | -0.0015 ($p_{\text{corr}} = .31$) |
| Group*Timepoint |  |  |  |  |
| CHR – Month 6 | -0.0020 ($p_{\text{corr}} = .24$) | -0.0024 ($p_{\text{corr}} = .24$) | -0.0022 ($p_{\text{corr}} = .24$) | -0.0013 ($p_{\text{corr}} = .43$) |
| FEP – Month 6 | 0.0016 ($p_{\text{corr}} = .81$) | 0.0062 ($p_{\text{corr}} = .81$) | 0.0079 ($p_{\text{corr}} = .81$) | 0.0027 ($p_{\text{corr}} = .81$) |
| CHR – Month 12 | 0.0013 ($p_{\text{corr}} = .51$) | 0.0012 ($p_{\text{corr}} = .51$) | 0.0011 ($p_{\text{corr}} = .51$) | 0.0016 ($p_{\text{corr}} = .51$) |
| FEP – Month 12 | 0.0014 ($p_{\text{corr}} = .65$) | 0.0008 ($p_{\text{corr}} = .65$) | 0.0010 ($p_{\text{corr}} = .65$) | 0.0014 ($p_{\text{corr}} = .65$) |

*Note.* Fixed-effect estimates (FDR-corrected $p$ -values) from linear mixed-effects model showed no significant effects of group, timepoint, and group-by-timepoint interactions on longitudinal changes in fractional anisotropy values. HC = Healthy Controls; CHR = Clinical High-Risk; FEP = First-Episode Psychosis; CB = Cingulum Bundle; SLF = Superior Longitudinal Fasciculus; IFOF = Inferior Fronto-Occipital Fasciculus.

**Figure 2.**
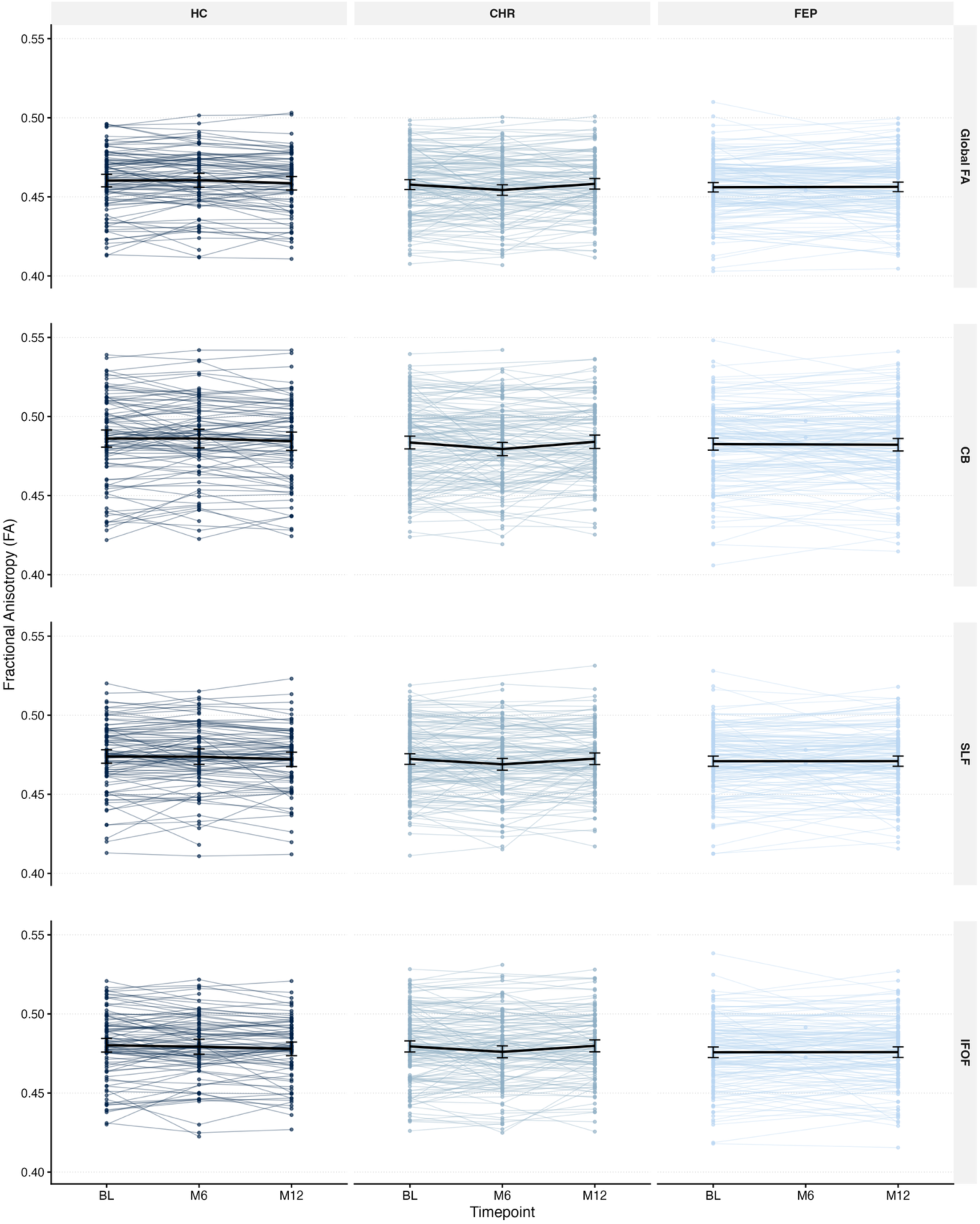
Longitudinal FA Trajectories by Group and Outcome. *Note.* FA trajectories in all regions of interest were not significantly associated with group status, time, or their interaction. HC = Healthy Controls; CHR = Clinical High-Risk; FEP = First-Episode Psychosis; CB = Cingulum Bundle; SLF = Superior Longitudinal Fasciculus; IFOF = Inferior Fronto-Occipital Fasciculus; BL = baseline; M6 = Month 6; M12 = Month 12.

In contrast, the dose of antipsychotic medication at baseline was significantly associated with lower FA across all tracts examined (global FA: *p_corr_* = .007; CB: *p_corr_* = .005; SLF: *p_corr_* = .02; and IFOF: *p_corr_* = .02; Tables S3-S6). For example, a 100 mg/day increase in Chlorpromazine equivalent was associated with a decrease of 0.002 in global FA across timepoints (95% CI [-0.0034, -0.0007]).

Ethnicity also showed robust associations with FA trajectories at both the global and tract-specific levels (global FA: *p_corr_* < .0001; CB: *p_corr_* < .0001; SLF: *p_corr_* < .0001; and IFOF: *p_corr_* < .0001). Follow-up fixed-effects estimates indicated that participants from East Asian backgrounds had consistently lower FA compared to those of African background (e.g., SLF: β = −0.0112, 95% CI [−0.0193, −0.0031]; Tables S3-S6).

No other covariates, including age, gender, or baseline IQ, showed significant associations with FA trajectories (*p_corr_* > .05; Table S3-S6).

### 2.3. Clinical Correlates of FA Trajectories

Consistent with the primary analysis, none of the examined clinical measures, (i) SOFAS, (ii) PANSS, or (iii) CAARMS, were significantly associated with FA trajectories either globally or within any tract (all *p_corr_* > .05; e.g. CB [SOFAS]: β = −0.0001, 95% CI [−0.0001, 0.0000], *p_corr_* = .69). Likewise, FA did not significantly differ depending on timepoint in any of the clinical models (all *p_corr_* > .05; e.g. SLF [SOFAS: Month 6 vs Baseline]: β = -0.0064, 95% CI [-0.0156, 0.0028]), or based on the interaction of time and clinical measure in any examined brain region (all *p_corr_* > .05; e.g. IFOF [CAARMS: CAARMS – Month 12 vs Baseline]: β = -0.0002, 95% CI [-0.0008, 0.0004], *p_corr_* = .53; Table S7-S9).

Baseline antipsychotic dose remained significantly associated with all FA trajectories in the SOFAS and PANSS models (e.g. SOFAS [global FA: *p_corr_* = .0005; CB: *p_corr_* = .0003; SLF: *p_corr_* = .007; and IFOF: *p_corr_* = .001]). For example, a 100 mg/day increase in Chlorpromazine equivalent was associated with a decrease of 0.002 in FA in the IFOF across timepoints within the SOFAS model (95% CI [-0.0035, -0.0009]), while a 100 mg/day increase was associated with a decrease of 0.003 in FA in the CB across timepoints within the PANSS model (95% CI [-0.0046, -0.0016]). Baseline antipsychotic dose was not significantly associated with FA in the CAARMS model (all *p_corr_* > .05), which only included CHR-P participants.

Ethnicity again emerged as a significant predictor of FA trajectories across models, showing robust associations with global FA as well as the CB, SLF, and IFOF in the SOFAS (all *p_corr_* < .0001) and CAARMS models (global FA: *p_corr_* = .005; CB: *p_corr_* = .005; SLF: *p_corr_* = .02; and IFOF: *p_corr_* = .005). Similar to the group-based analysis, this effect was driven by participants from East Asian backgrounds, who consistently exhibited lower FA relative to other ethnic groups (e.g., global FA [SOFAS]: β = -0.0136, 95% CI [-0.0293, -0.0103], *p_corr_* = .0005; SLF [CAARMS]: β = -0.0139, 95% CI [-0.0268, -0.0010], *p_corr_* = .04). This pattern was not observed in the PANSS model (all *p_corr_* > .05).

Finally, as in the primary models, the covariates of age, gender, and baseline IQ did not emerge as significant predictors of FA change (*p_corr_* > .05).

### 2.4. Sensitivity Analyses

The pattern observed in the hemisphere-specific analyses (a), gender-stratified analyses (b), and models excluding baseline antipsychotic dose (c) closely paralleled those of the primary models. Neither the main effects of group status and time, nor the group-by-time interaction, were significantly associated with changes in FA across investigated areas (all *p_corr_* > .05; Table S10-S14). A closer inspection of post hoc model-based contrasts revealed a marginally significant reduction in FA trajectories in the FEP group compared with healthy controls within the CB (β = -0.0068, 95% CI [-0.0138, 0.0002], *p_corr_* = .076), the IFOF (β = -0.0064, 95% CI [-0.0124, -0.0004], *p_corr_* = .072), and globally (β = -0.0060, 95% CI [-0.0114, -0.0006], *p_corr_* = .072), when baseline antipsychotic dose was excluded from the model.

Higher baseline antipsychotic doses were significantly associated with greater longitudinal decline in FA. This effect was observed across all WM tracts except the left SLF in the hemisphere-specific models (*p_corr_* < .05), and globally as well as across all tracts except the SLF in men in the gender-stratified models (*p_corr_* < .05). For example, an increase of 100 mg/day was associated with a decrease of 0.003 in FA in the right CB across timepoints (95% CI [-0.0048, -0.0013]). Notably, no significant association between antipsychotic dose and FA change was observed in women (model (b); *p_corr_* > .05).

Again, ethnicity remained the most consistent predictor of changes in FA after correction for multiple comparisons. In all sensitivity analyses models, East Asian ethnicity was significantly associated with greater FA decreases both globally and across all tracts in the hemisphere-specific (*p_corr_* < .001), antipsychotic-free models (*p_corr_* < .001), and gender-stratified (all *p_corr_* < .01; e.g. IFOF in women: β = -0.0075, 95% CI [-0.0383, -0.0087], *p_corr_* = .005).

Age, gender, and baseline IQ were not significant predictors of FA change across any of the sensitivity analyses (*p_corr_* > .05).

### 2.5. Post Hoc Analyses

Antipsychotic dose at baseline was negatively correlated with SOFAS scores across groups (*r* = -.29, *p* < .001) and positively correlated with PANSS positive scores within the FEP group (*r* = .22, *p* = .006; Figures S1-S2). In the gender-stratified clinical scores comparison, males showed significantly higher baseline antipsychotic doses (*p* = .003), descriptively higher PANSS positive scores (*p* = .14), and marginally lower SOFAS scores (*p* = .06), compared to female participants within the FEP group (Table S15).

## 3. Discussion

This study examined longitudinal changes in WM microstructure during the clinical-high-risk and first-episode phases of psychosis. Contrary to our first hypothesis, there were no differences between people at CHR-P, patients with FEP, and HC in FA trajectories over 12 months, either in the selected WM tracts or at the global level. In addition, we did not find relationships between longitudinal changes in FA and (attenuated) psychotic symptom severity or level of functioning. However, we found that lower levels of FA were related to higher doses of antipsychotic medication at baseline, and to East Asian ethnicity.

The absence of robust group differences in longitudinal FA suggests that there are no consistent or clinically meaningful differences in WM changes over a 12-month period between healthy individuals and those with CHR-P or FEP status. Because this null effect persisted across all model specifications, it is unlikely to be driven solely by insufficient statistical power. Instead, the findings indicate that any true group differences in FA trajectory are likely small in magnitude and/or highly heterogeneous. The consistency of null findings in global FA and all analysed tracts further suggests that these results reflect widespread overlap in white-matter microstructural trajectories across groups, rather than region-specific effects. Moreover, the absence of an interaction of group status and time argues against a model of rapid or accelerated WM degeneration during CHR-P and FEP over the observed period.

Importantly, these findings help contextualise the heterogeneous longitudinal patterns reported in the existing literature. While some studies and reviews have described increasing FA over time in CHR-P samples ^14^, and others have reported largely stable or declining trajectories in similar populations ^12^, our results suggest that such discrepancies may reflect substantial inter-individual variability, short follow-up periods, and sensitivity to methodological choices rather than robust, diagnosis-specific trajectories. In this context, the lack of consistent group differences observed here is not inconsistent with prior work, but rather supports the view that longitudinal FA changes in CHR-P and FEP are highly variable and weakly constrained by categorical diagnostic status. Notably, we observed regional FA differences between FEP and HC only in models that did not account for baseline antipsychotic exposure, underscoring the critical importance of rigorous covariate control when interpreting longitudinal DTI findings. Together, these results suggest that diagnostic categories alone may be poor proxies for short-term neurobiological change, and that observed longitudinal variation in FA is more likely to reflect developmental, demographic, and treatment-related factors rather than core diagnostic processes per se. The apparent relative stability of FA over the observed interval suggests that, at least over short time frames, cross-sectional FA measures may be less sensitive to the precise timing of image acquisition, cautiously supporting the use of cross-sectional differences in FA as a biomarker ^16^.

Longitudinal functional impairment and psychotic symptom severity were not robustly associated with FA trajectories. The consistency of null findings across clinical instruments argues against instrument-specific insensitivity and instead points towards a conceptual decoupling between clinical expression and changes in WM microstructure over the observed period. One possible explanation is that these processes operate on different temporal scales: changes in white-matter microstructure, as assessed by DTI, may evolve gradually, whereas clinical symptoms fluctuate more dynamically over shorter intervals ^48^. In addition, microstructural alterations may operate upstream of the functional network disruptions that give rise to clinical symptoms, with effects that are indirect and temporally lagged. Such asynchrony between biological and clinical processes may therefore attenuate observable associations in longitudinal analyses. This interpretation is consistent with models in which white-matter alterations reflect trait-like vulnerability or neurodevelopmental factors, while clinical symptoms are more closely linked to dynamic network-level and neurochemical processes ^48,49^.

Baseline antipsychotic dose emerged as one of the most reproducible correlates of FA decline, primarily in FEP. The consistency of this association across tracts suggests a global or diffuse influence rather than a tract-specific vulnerability, and aligns with direct medication effects on brain structure and indirect effects via illness severity or treatment duration ^50–52^. Importantly, a higher baseline antipsychotic dose was associated with lower levels of functioning across all groups and with greater positive symptom severity within FEP. In addition, male FEP participants showed higher levels of antipsychotic dose and lower levels of functioning compared to females. Together, these findings indicate that antipsychotic dose may act as a proxy for illness severity, which could also account for the observed association with FA decline, as well as the absence of medication effects in females. One possible explanation for the stronger association of antipsychotic dose compared to clinical measures in our study is that medication exposure prior to study enrolment may have had beneficial effects on symptom severity, thereby weakening associations between clinical scales and FA, whilst preserving the relationship with antipsychotic dose. Notably, previous studies have also reported positive associations between antipsychotic exposure and FA increases ^53,54^. Together with our findings, this suggests that antipsychotic-related WM changes may reflect a combination of therapeutic, compensatory, and illness-related processes. Importantly, causality or directionality of the association between antipsychotic exposure and WM microstructure cannot be inferred due to the design of the study, and this finding should be treated with care.

Ethnicity was consistently associated with longitudinal FA across analytic models, suggesting it may reflect an important contextual factor. This association may have been driven by unmeasured factors, such as environmental exposures, socioeconomic conditions, and differences in healthcare systems and early detection pathways ^55^. Similarly, despite extensive harmonisation, residual site effects may also have contributed to this finding. Taken together with the significance of antipsychotic exposure in our models, these results caution against interpreting group or clinical differences in WM measures without careful control for demographic and medication-related factors, as residual confounding may obscure or inflate disease-related effects. Accordingly, explicit modelling of antipsychotic dose should be prioritised in neuroimaging studies, particularly in early psychosis, where dosing varies widely.

### 3.1. Limitations and Future Directions

While the longitudinal, multi-centre design of this study represents a clear strength compared to cross-sectional and single-site studies, it also entails limitations. To ensure feasibility across sites, DTI acquisition protocols were constrained, resulting in single-shell DTI data. Consequently, more advanced DTI analysis approaches that aim to overcome the interpretational limits of FA, such as NODDI ^56^ or BENCH ^57^, could not be applied. Although the cohort includes multiple countries, group sizes for some ethnicities were small, and global and ethnic representativeness could be further improved through greater inclusion of underrepresented regions and groups. In addition, the 12-month imaging interval may be insufficient to capture slower-evolving white-matter changes, and antipsychotic exposure was only available at baseline, limiting longitudinal medication analysis. Future studies should aim to build on the strengths of this multi-centre longitudinal framework by extending follow-up periods, improving global representativeness, and harmonising acquisition protocols in a way that enables the application of more advanced diffusion models.

From an analytic perspective, our focus on three major tracts was guided by prior literature. However, given the inconsistency of existing findings, relevant effects may exist in other WM pathways not examined here, highlighting the potential value of more data-driven approaches to detect distributed patterns of WM change. Similarly, to better capture the heterogeneity, future work should examine clinically meaningful subgroups, including individuals at CHR-P who transition to psychosis versus those who do not and treatment-resistant FEP. Detecting such transition-related effects will likely require analytic approaches specifically designed for this purpose, as the present framework is underpowered for these subgroup comparisons.

### 3.2. Conclusions

In this prospective, multi-centre longitudinal study, we examined trajectories of WM microstructure during CHR-P and FEP. Contrary to our hypotheses, we found no evidence for differential longitudinal changes in FA between healthy controls, individuals at clinical high risk for psychosis, and those experiencing a first episode of psychosis, either globally or within key association tracts. WM microstructure was also not robustly associated with symptom severity or functionality. However, there was evidence that the longitudinal trajectory of FA is related to the dose of antipsychotic medication at baseline. These findings suggest that large-scale WM microstructural changes during early psychosis are subtle and heterogeneous. Future studies with longer follow-up, more sensitive imaging and analytic approaches, and careful consideration of covariates will be needed to clarify the mechanisms underlying WM alterations in psychosis.

## Acknowledgements

Special thanks to Gaurav Bhalerao and Jacob Turnbull for their expertise in site harmonisation and Jesper Anderson for his guidance during the preprocessing of the diffusion data.

## 6. Collaborators

Alexis E. Cullen ^1^, Kate Merritt ^1^, Andrea Mechelli ^1^, Natalia Petros ^1^, Mathilde Antoniades ^1^, Andrea De Micheli ^1^, Sandra Vieira ^1^, Tom Spencer ^1^, Gemma Modinos ^2^, Frederike Schirmbeck ^3^, Diana Tordesillas-Gutierrez ^4,5^, Esther Setien-Suero ^4,5^, Rosa Avesa-Arriola ^4,5^, Paula Suarez-Pinilla ^4,5^, Victor Ortiz Garcia-de la Foz ^4,5^, Mikkel Erlang Sørensen ^6^, Bjørn H. Ebdrup ^6,7^, Karen Tangmose ^6,7^, Helle Schæbel ^6^, Egill Rostrup ^6,8^, Brian Hallahan ^9^, Dara Cannon ^9^, James McLoughlin ^9^, Martha Finnegan ^9^, Oliver Gruber ^10^, Anja Richter ^1,10^, Bernd Krämer ^10^, Bea Campforts ^11^, Machteld Marcelis ^11^, Claudia Vingerhoets ^11^, Covandonga M. Díaz-Caneja ^12^, Miriam Avora ^12^, Joost Janssen ^12^, Roberto Rodríguez-Jiménez ^13^, Marina Díaz-Marsá ^14^, Tilo Kircher ^15^, Florian Bitsch ^15^, Jens Sommer ^15^, Patrick McGorry ^16,17^, Paul Amminger ^16,17^, Meredith McHugh ^16,17^, Armida Mucci ^18^, Paola Bucci ^18^, Giuseppe Piegari ^18^, Daria Pietrafesa ^18^, Alessia Nicita ^18^, Sara Patriarca ^18^, André Zugman ^19^, Graccielle Rodrigues da Cunha ^19^, Tae Young Lee ^20^, Minah Kim ^20^, Sun-Young Moon ^20^, Silvia Kyungjin Lho ^20^, Michael Kiang ^21,22,23^, Sarah Ahmed ^21,22^, Jenny Lepock ^21,22^, Margaret Maheandiran ^22^, Ivana Prce ^22^, Cory Gerritsen ^22,24^, Matthäus Willeit ^25^, Marzena Lenczowski ^25^, Ulrich Sauerzopf ^25^, Ana Weidenauer ^25^, Julia Furtner-Srajer ^26^, Anke Maatz ^27^, Achim Burrer ^27^, Philipp Stämpfli ^27^, Naemi Huber ^27^, Stefan Kaiser ^28,29^, Wolfram Kawohl ^30^

1. Department of Psychosis Studies, Institute of Psychiatry, Psychology & Neuroscience, King’s College London, London, UK
2. Department of Psychological Medicine, Institute of Psychiatry, Psychology & Neuroscience, King’s College London, London, UK
3. Amsterdam UMC, University of Amsterdam, Psychiatry, Department Early Psychosis, Amsterdam, The Netherlands
4. Department of Psychiatry, Marqués de Valdecilla University Hospital, IDIVAL. School of Medicine, University of Cantabria, Santander, Spain
5. CIBERSAM, Centro Investigación Biomédica en Red Salud Mental, Spain
6. Centre for Neuropsychiatric Schizophrenia Research (CNSR) & Centre for Clinical Intervention and Neuropsychiatric Schizophrenia Research (CINS), Mental Health Centre Glostrup, University of Copenhagen, Glostrup, Denmark
7. University of Copenhagen, Faculty of Health and Medical Sciences, Deptartment of Clinical Medicine, Copenhagen, Denmark
8. Functional Imaging Unit (FIUNIT), Rigshospitalet Glostrup, University of Copenhagen, Glostrup, Denmark
9. Centre for Neuroimaging, Cognition and Genomics (NICOG), Galway Neuroscience Centre, College of Medicine Nursing and Health Sciences, University of Galway, Galway, Ireland
10. Section for Experimental Psychopathology and Neuroimaging, Department of General Psychiatry, Heidelberg University, Heidelberg, Germany
11. Department of Psychiatry and Neuropsychology, Maastricht University, Maastricht, The Netherlands
12. Servicio de Psiquiatría del Niño y del Adolescente, Hospital General Universitario Gregorio Marañon, Universidad Complutense Madrid, Spain; Centro de Investigación Biomédica en Red de Salud Mental, Madrid, Spain
13. Departmento de Psiquiatría, Instituto de Investigación Sanitaria Hospital 12 de Octubre (imas12), Madrid, Spain
14. Hospital Clínico de San Carlos, Universidad Complutense, Centro de Investigación Biomédica en Red de Salud Mental (CIBERSAM), Madrid, Spain
15. Department of Psychiatry, University of Marburg, Marburg, Germany
16. Orygen, 35 Poplar Road, Parkville, Victoria, Melbourne, Australia
17. Centre for Youth Mental Health, The University of Melbourne, Parkville, Victoria, Australia
18. Department of Psychiatry, University of Campania Luigi Vanvitelli, Largo Madonna delle Grazie, Naples, Italy
19. Department of Psychiatry, Interdisciplinary Lab for Clinical Neurosciences (LiNC), Universidade Federal de São Paulo (UNIFESP), São Paulo, Brazil
20. Department of Psychiatry, Seoul National University College of Medicine, Jongno-gu, Seoul, South Korea
21. Institute of Medical Science, University of Toronto, Toronto, Ontario, Canada
22. Centre for Addiction and Mental Health, Toronto, Ontario, Canada
23. Department of Psychiatry, McGill University, Montreal, Canada
24. Department of Psychology, University of Toronto, Toronto, Ontario, Canada
25. Department of Psychiatry and Psychotherapy, Medical University of Vienna, Vienna, Austria
26. Department of Biomedical Imaging and Image-guided Therapy, Medical University of Vienna, Vienna, Austria
27. Department of Psychiatry, Psychotherapy and Psychosomatics, Psychiatric Hospital, University of Zurich, Switzerland
28. Clinical and Experimental Psychopathology Laboratory, Faculty of Medicine, University of Geneva, Switzerland
29. Adult Psychiatry Division, Department of mental health and psychiatry, University Hospitals of Geneva, Switzerland
30. Department for Psychiatry and Psychotherapy, Psychiatric Services Aargau, Brugg, Switzerland

