## Supplementary material for "Longitudinal White Matter Trajectories in Clinical High Risk and First-Episode Psychosis: Findings from the Multi-Centre PSYSCAN Study": Table S1

*Acquisition Parameters for Diffusion-Weighted Magnetic Resonance Images per PSYSCAN Site*

| Site | Scanner Make | Scanner Model | Coil[a] | Repetition Time | Echo Time | Pixel Spacing | Slices | Gradient Directions [b] | b0 Volumes |
|---|---|---|---|---|---|---|---|---|---|
| Melbourne, AU | Siemens | MAGNETOM Prisma$^{Fit}$ | 64 | 7900 | 88 | 2.5 x 2.5 | 64 | 64 | 5 |
| Vienna, AT | Siemens | MAGNETOM TrioTim | 12 | 8100 | 88 | 2.5 x 2.5 | 64 | 64 | 1 |
| Copenhagen, DK | Philips | Achieva dStream | 32 | 6805 | 74 | 2.5 x 2.5 | 64 | 64 | 1 |
| Marburg, DE | Siemens | MAGNETOM TrioTim | 12 | 7800 | 88 | 2.5 x 2.5 | 64 | 64 | 5 |
| Galway, IE | Philips | Achieva TX | 32 | 6824 | 74 | 2.5 x 2.5 | 64 | 64 | 1 |
| Tel Hashomer, IL | GE | Signa HDxT | 8 | 14600 | 84.6 | 1.25 x 1.25 | 64 | 64 | 5 |
| Naples, IT | Siemens | MAGNETOM TrioTim | 8 | 7900 | 89 | 2.5 x 2.5 | 64 | 64 | 5 |
| Amsterdam, NL | Philips | Ingenia | 32 | 7259.37 | 82 | 2.5 x 2.5 | 64 | 64 | 1 |
| Maastricht, NL | Siemens | MAGNETOM Prisma$^{Fit}$ | 64 | 7700 | 88 | 2.5 x 2.5 | 64 | 64 | 5 |
| Utrecht, NL | Philips | Ingenia CX | 32 | 65601 | 74 | 2.5 x 2.5 | 64 | 64 | 1 |
| Cantabria, ES | Philips | Achieva | 8 | 6559.26 | 74 | 2.5 x 2.5 | 64 | 64 | 1 |
| Madrid, ES | GE | Signa HDxT | 8 | 13100 | 69.4 | 2.5 x 2.5 | 62 | 64 | 5 |
| Zurich, CH | Philips | Achieva dStream | 32 | 6559.8 | 74 | 2.5 x 2.5 | 64 | 64 | 1 |
| Edinburgh, UK | Siemens | MAGNETOM Verio | 12 | 9800 | 88 | 2.5 x 2.5 | 64 | 64 | 5 |
| London, UK | GE | Discovery MR750 | 12 | 10000 | 68.3 | 2.5 x 2.5 | 64 | 64 | 5 |
| Toronto, CA | GE | Discovery MR750 | 8 | 10000 | 68.3 | 2.5 x 2.5 | 64 | 64 | 5 |
| Seoul, KR | Siemens | MAGNETOM TrioTim | 12 | 7800 | 88 | 2.5 x 2.5 | 64 | 64 | 5 |
| Sao Paulo, BR | Philips | Achieva | 32 | 6479 | 74 | 2.5 x 2.5 | 64 | 32 | 1 |

*Note.* The orientation was axial for all sites

[a] Number of channels. [b] b = 1000 $m/s^2$

**Table S2**

*Conversion of CAARMS and PANSS into SOPS*

| SOPS | PANSS | | CAARMS | |
|---|---|---|---|---|
| Item | Item | PANSS to SOPS Scoring | Item | CAARMS to SOPS Scoring |
| P1 – Unusual Thought Content / Delusional Ideas | P1 – Delusions | 1 = 0 | Unusual Thought Content | 0 = 0.012 |
| | | 2 = 3 | | 1 = 1.092 |
| | | 3 = 4 | | 2 = 2.212 |
| | | 4 = 6 | | 3 = 3.258 |
| | | 5 = 6 | | 4 = 4.180 |
| | | 6 = 7 | | 5 = 5.019 |
| | | 7 = 7 | | 6 = 5.965 |
| P2 – Suspiciousness / Persecutory Ideas | P6 – Suspiciousness / Persecution | 1 = 0 | Non-Bizarre Ideas | 0 = 0.026 |
| | | 2 = 3 | | 1 = 1.059 |
| | | 3 = 4 | | 2 = 2.234 |
| | | 4 = 6 | | 3 = 3.216 |
| | | 5 = 6 | | 4 = 4.105 |
| | | 6 = 7 | | 5 = 5.956 |
| | | 7 = 7 | | 6 = 5.961 |
| P4 – Perceptual Abnormalities / Hallucinations | P3 – Hallucinatory Behaviour | 1 = 0 | Perceptual Abnormalities | 0 = 0.012 |
| | | 2 = 3 | | 1 = 0.893 |
| | | 3 = 4 | | 2 = 1.891 |
| | | 4 = 6 | | 3 = 2.953 |
| | | 5 = 6 | | 4 = 3.944 |
| | | 6 = 7 | | 5 = 4.861 |

| | | | | |
|---|---|---|---|---|
| | | 7 = 7 | | 6 = 5.876 |
| P5 – Disorganized Communication | P2 – Conceptual Disorganisation | 1 = 0 | Disorganised Speech | 0 = 0.067 |
| | | 2 = 3 | | 1 = 0.855 |
| | | 3 = 4 | | 2 = 1.981 |
| | | 4 = 6 | | 3 = 3.032 |
| | | 5 = 6 | | 4 = 4.064 |
| | | 6 = 7 | | 5 = 5.158 |
| | | 7 = 7 | | 6 = 6.090 |

*Note.* Based on the works of Fusar-Poli et al. (2016) and Schobel et al. (2009).

**Table S3**

*Primary Analysis Fixed Effects – Global FA*

| Fixed Effect | est | *SE* | 95 % CI | *t/F* | *p* | *p* corr |
|---|---|---|---|---|---|---|
| Group | | | | 0.4791 | .6196 | .8766 |
| CHR vs HC | -0.0021 | 0.0025 | [-0.0069, 0.0028] | -0.8276 | .4084 | .7224 |
| FEP vs HC | -0.0020 | 0.0031 | [-0.0081, 0.0041] | -0.6307 | .5286 | .9184 |
| Timepoint | | | | 0.4607 | .6311 | .7373 |
| Month 6 | 0.0003 | 0.0011 | [-0.0019, 0.0025] | 0.2618 | .7936 | .9016 |
| Month 12 | -0.0014 | 0.0011 | [-0.0037, 0.0008] | -1.2746 | .2030 | .3130 |
| Age | 0.0002 | 0.0002 | [-0.0001, 0.0006] | 1.3747 | .1701 | .3401 |
| Gender | -0.0041 | 0.0020 | [-0.0080, -0.0002] | -2.0834 | *.0379 | .0758 |
| Ethnicity | | | | 7.0500 | *<.0001 | *<.0001 |
| Asian Indian vs African | -0.0032 | 0.0057 | [-0.0145, 0.0081] | -0.5576 | .5775 | .7699 |
| East Asian vs African | -0.0137 | 0.0038 | [-0.0211, -0.0063] | -3.6371 | *.0003 | *.0004 |
| Inter Racial vs African | -0.0169 | 0.0107 | [-0.0379, 0.0041] | -1.5840 | .1141 | .1638 |
| Middle Eastern vs African | 0.0125 | 0.0083 | [-0.0038, 0.0287] | 1.5054 | .1331 | .3695 |
| Other vs African | -0.0041 | 0.0046 | [-0.0131, 0.0048] | -0.9064 | .3653 | .4871 |
| White vs African | 0.0037 | 0.0028 | [-0.0018, 0.0093] | 1.3144 | .1895 | .3790 |
| CPZ (mg/day) | 0.0000 | 0.0000 | [0, 0] | -2.9191 | *.0037 | *.0074 |
| WAIS | 0.0001 | 0.0001 | [0, 0.0002] | 1.0669 | .2867 | .4663 |
| Group x Timepoint | | | | 1.2052 | .3076 | .4446 |
| CHR – Month 6 | -0.0020 | 0.0015 | [-0.0048, 0.0009] | -1.3519 | .1770 | .2367 |
| FEP – Month 6 | 0.0016 | 0.0065 | [-0.0113, 0.0144] | 0.2414 | .8094 | .8094 |
| CHR – Month 12 | 0.0013 | 0.0015 | [-0.0016, 0.0042] | 0.8747 | .3822 | .5133 |
| FEP – Month 12 | 0.0014 | 0.0014 | [-0.0014, 0.0041] | 0.9841 | .3255 | .6487 |

*Note.* Fixed effects from the primary analysis linear mixed-effects model (random intercept for subject) for global FA values; HC and baseline are reference categories. P values are FDR-corrected across primary models.

* $p < .05$

**Table S4**

*Primary Analysis Fixed Effects - CB*

| Fixed Effect | est | *SE* | 95 % CI | *t/F* | *p* | *p* corr |
|---|---|---|---|---|---|---|
| Group | | | | 0.4197 | .6575 | .8766 |
| CHR vs HC | -0.0016 | 0.0032 | [-0.0079, 0.0047] | -0.4989 | .6181 | .7224 |
| FEP vs HC | -0.0010 | 0.0040 | [-0.0088, 0.0069] | -0.2459 | .8059 | .9184 |
| Timepoint | | | | 0.7808 | .4585 | .7373 |
| Month 6 | 0.0002 | 0.0014 | [-0.0025, 0.0029] | 0.1237 | .9016 | .9016 |
| Month 12 | -0.0014 | 0.0014 | [-0.0041, 0.0013] | -1.0304 | .3033 | .3130 |
| Age | 0.0004 | 0.0002 | [-0.0001, 0.0008] | 1.6637 | .0970 | .3401 |
| Gender | -0.0060 | 0.0026 | [-0.0111, -0.0010] | -2.3580 | *.0189 | .0756 |
| Ethnicity | | | | 7.4334 | *<.0001 | *<.0001 |
| Asian Indian vs African | -0.0063 | 0.0074 | [-0.0209, 0.0083] | -0.8512 | .3952 | .7699 |
| East Asian vs African | -0.0198 | 0.0049 | [-0.0293, -0.0102] | -4.0517 | *.0001 | *.0001 |
| Inter Racial vs African | -0.0213 | 0.0138 | [-0.0484, 0.0058] | -1.5466 | .1228 | .1638 |
| Middle Eastern vs African | 0.0116 | 0.0107 | [-0.0094, 0.0327] | 1.0885 | .2771 | .3695 |
| Other vs African | -0.0063 | 0.0059 | [-0.0179, 0.0054] | -1.0599 | .2899 | .4871 |
| White vs African | 0.0039 | 0.0036 | [-0.0033, 0.0111] | 1.0694 | .2856 | .3808 |
| CPZ (mg/day) | 0.0000 | 0.0000 | [0, 0] | -3.2415 | *.0013 | *.0052 |
| WAIS | 0.0000 | 0.0001 | [-0.0001, 0.0002] | 0.6981 | .4856 | .4856 |
| Group x Timepoint | | | | 1.1472 | .3334 | .4446 |
| CHR – Month 6 | -0.0024 | 0.0018 | [-0.0060, 0.0011] | -1.3648 | .1729 | .2367 |
| FEP – Month 6 | 0.0062 | 0.0081 | [-0.0096, 0.0220] | 0.7687 | .4424 | .8094 |
| CHR – Month 12 | 0.0012 | 0.0018 | [-0.0024, 0.0047] | 0.6542 | .5133 | .5133 |
| FEP – Month 12 | 0.0008 | 0.0017 | [-0.0026, 0.0041] | 0.4558 | .6487 | .6487 |

*Note.* Fixed effects from the primary analysis linear mixed-effects model (random intercept for subject) for the cingulum bundle.

* $p < .05$

**Table S5**

*Primary Analysis Fixed Effects - SLF*

| Fixed Effect | est | *SE* | 95 % CI | *t/F* | *p* | *p* corr |
|---|---|---|---|---|---|---|
| Group | | | | 0.6150 | .5410 | .8766 |
| CHR vs HC | -0.0010 | 0.0027 | [-0.0064, 0.0043] | -0.3806 | .7037 | .7224 |
| FEP vs HC | -0.0003 | 0.0034 | [-0.0070, 0.0063] | -0.1025 | .9184 | .9184 |
| Timepoint | | | | 0.8821 | .4144 | .7373 |
| Month 6 | 0.0003 | 0.0013 | [-0.0021, 0.0028] | 0.2693 | .7878 | .9016 |
| Month 12 | -0.0013 | 0.0013 | [-0.0038, 0.0012] | -1.0098 | .3130 | .3130 |
| Age | 0.0002 | 0.0002 | [-0.0002, 0.0006] | 0.8816 | .3786 | .4008 |
| Gender | -0.0032 | 0.0022 | [-0.0074, 0.0011] | -1.4645 | .1439 | .1919 |
| Ethnicity | | | | 5.7448 | *<.0001 | *<.0001 |
| Asian Indian vs African | 0.0011 | 0.0063 | [-0.0112, 0.0135] | 0.1810 | .8565 | .8565 |
| East Asian vs African | -0.0112 | 0.0041 | [-0.0193, -0.0031] | -2.7310 | *.0066 | *.0066 |
| Inter Racial vs African | -0.0119 | 0.0116 | [-0.0348, 0.0110] | -1.0225 | .3072 | .3072 |
| Middle Eastern vs African | 0.0103 | 0.0090 | [-0.0075, 0.0281] | 1.1397 | .2552 | .3695 |
| Other vs African | -0.0007 | 0.0050 | [-0.0106, 0.0091] | -0.1500 | .8809 | .8809 |
| White vs African | 0.0063 | 0.0031 | [0.0003, 0.0124] | 2.0519 | *.0409 | .1635 |
| CPZ (mg/day) | 0.0000 | 0.0000 | [0, 0] | -2.2648 | *.0241 | *.0241 |
| WAIS | 0.0001 | 0.0001 | [0, 0.0002] | 1.2686 | .2054 | .4663 |
| Group x Timepoint | | | | 1.3292 | .2578 | .4446 |
| CHR – Month 6 | -0.0022 | 0.0016 | [-0.0054, 0.0010] | -1.3502 | .1775 | .2367 |
| FEP – Month 6 | 0.0079 | 0.0073 | [-0.0065, 0.0223] | 1.0744 | .2831 | .8094 |
| CHR – Month 12 | 0.0011 | 0.0016 | [-0.0021, 0.0043] | 0.6701 | .5031 | .5133 |
| FEP – Month 12 | 0.0010 | 0.0016 | [-0.0020, 0.0041] | 0.6472 | .5178 | .6487 |

*Note.* Fixed effects from the primary analysis linear mixed-effects model (random intercept for subject) for the superior longitudinal fasciculus.

* $p < .05$

**Table S6**

*Primary Analysis Fixed Effects – IFOF*

| Fixed Effect | est | *SE* | 95 % CI | *t/F* | *p* | *p* corr |
|---|---|---|---|---|---|---|
| Group | | | | 0.0761 | .9267 | .9267 |
| CHR vs HC | -0.0010 | 0.0027 | [-0.0064, 0.0044] | -0.3555 | .7224 | .7224 |
| FEP vs HC | -0.0027 | 0.0034 | [-0.0095, 0.0040] | -0.7918 | .4289 | .9184 |
| Timepoint | | | | 0.3049 | .7373 | .7373 |
| Month 6 | -0.0005 | 0.0012 | [-0.0029, 0.0019] | -0.4059 | .6850 | .9016 |
| Month 12 | -0.0015 | 0.0012 | [-0.004, 0.0009] | -1.2238 | .2215 | .3130 |
| Age | 0.0002 | 0.0002 | [-0.0002, 0.0006] | 0.8411 | .4008 | .4008 |
| Gender | -0.0026 | 0.0022 | [-0.0069, 0.0018] | -1.1642 | .2451 | .2451 |
| Ethnicity | | | | 6.7047 | *<.0001 | *<.0001 |
| Asian Indian vs African | -0.0056 | 0.0064 | [-0.0181, 0.0069] | -0.8868 | .3758 | .7699 |
| East Asian vs African | -0.0188 | 0.0042 | [-0.0270, -0.0105] | -4.4870 | *<.0001 | *<.0001 |
| Inter Racial vs African | -0.0248 | 0.0118 | [-0.0481, -0.0016] | -2.1020 | *.0362 | .1449 |
| Middle Eastern vs African | 0.0071 | 0.0092 | [-0.0110, 0.0251] | 0.7704 | .4415 | .4415 |
| Other vs African | -0.0066 | 0.0051 | [-0.0165, 0.0034] | -1.2958 | .1958 | .4871 |
| White vs African | 0.0002 | 0.0031 | [-0.0060, 0.0064] | 0.0635 | .9494 | .9494 |
| CPZ (mg/day) | 0.0000 | 0.0000 | [0, 0] | -2.4052 | *.0166 | *.0222 |
| WAIS | 0.0001 | 0.0001 | [-0.0001, 0.0002] | 0.9364 | .3497 | .4663 |
| Group x Timepoint | | | | 0.7911 | .5312 | .5312 |
| CHR – Month 6 | -0.0013 | 0.0016 | [-0.0044, 0.0019] | -0.7918 | .4289 | .4289 |
| FEP – Month 6 | 0.0027 | 0.0072 | [-0.0115, 0.0169] | 0.3716 | .7103 | .8094 |
| CHR – Month 12 | 0.0016 | 0.0016 | [-0.0015, 0.0048] | 1.0055 | .3151 | .5133 |
| FEP – Month 12 | 0.0014 | 0.0015 | [-0.0016, 0.0044] | 0.9330 | .3512 | .6487 |

*Note.* Fixed effects from the primary analysis linear mixed-effects model (random intercept for subject) for the inferior fronto-occipital fasciculus.

* $p < .05$

**Table S7**

*Secondary Analysis Fixed Effects – SOFAS*

| Fixed Effect | est | 95 % CI | *p* | *p* corr |
|---|---|---|---|---|
| | | Global FA | | |
| SOFAS | -0.0000 | [-0.0001, 0.0000] | .5778 | .6916 |
| Timepoint | | | | |
| Month 6 | -0.0038 | [-0.0120, 0.0044] | .3673 | .3673 |
| Month 12 | -0.0001 | [-0.0046, 0.0044] | .9554 | .9554 |
| SOFAS x Timepoint | | | | |
| SOFAS – Month 6 | 0.0000 | [-0.0001, 0.0002] | .5854 | .5854 |
| SOFAS – Month 12 | 0.0000 | [-0.0001, 0.0001] | .9799 | .9799 |
| | | Cingulum Bundle | | |
| SOFAS | -0.0001 | [-0.0001, 0.0000] | .2208 | .6916 |
| Timepoint | | | | |
| Month 6 | -0.0064 | [-0.0165, 0.0037] | .2150 | .3386 |
| Month 12 | -0.0005 | [-0.0060, 0.0051] | .8702 | .9554 |
| SOFAS x Timepoint | | | | |
| SOFAS – Month 6 | 0.0001 | [-0.0001, 0.0002] | .3841 | .5723 |
| SOFAS – Month 12 | 0.0000 | [-0.0001, 0.0001] | .9106 | .9799 |
| | | Superior Longitudinal Fasciculus | | |
| SOFAS | -0.0000 | [-0.0001, 0.0001] | .6104 | .6916 |
| Timepoint | | | | |
| Month 6 | -0.0064 | [-0.0156, 0.0028] | .1746 | .3386 |
| Month 12 | -0.0008 | [-0.0059, 0.0042] | .7496 | .9554 |
| SOFAS x Timepoint | | | | |
| SOFAS – Month 6 | 0.0001 | [-0.0001, 0.0002] | .2994 | .5723 |
| SOFAS – Month 12 | 0.0000 | [-0.0001, 0.0001] | .7709 | .9799 |
| | | Inferior Fronto-Occipital Fasciculus | | |
| SOFAS | -0.0000 | [-0.0001, 0.0001] | .6916 | .6916 |
| Timepoint | | | | |
| Month 6 | -0.0053 | [-0.0144, 0.0038] | .2539 | .3386 |
| Month 12 | -0.0014 | [-0.0064, 0.0036] | .5874 | .9554 |
| SOFAS x Timepoint | | | | |
| SOFAS – Month 6 | 0.0001 | [-0.0001, 0.0002] | .4292 | .5723 |
| SOFAS – Month 12 | 0.0000 | [-0.0001, 0.0001] | .6060 | .9799 |

*Note.* Fixed effects from the secondary analysis linear mixed-effects model (random intercept for subject); testing for effects of SOFAS and time on fractional anisotropy (FA), while controlling for age, gender, ethnicity, baseline antipsychotic dose, and baseline IQ. Benjamin-Hochberg correction was applied to adjust for multiple comparisons. None of the outcomes of interest (SOFAS, timepoint, SOFAS x Timepoint) reached statistical significance.

**Table S8**

*Secondary Analysis Fixed Effects – PANSS*

| Fixed Effect | est | 95 % CI | *p* | *p* corr |
|---|---|---|---|---|
| | | Global FA | | |
| PANSS | 0.0003 | [-0.0001, 0.0006] | .0977 | .1303 |
| Timepoint | 0.0036 | [-0.0018, 0.0090] | .1968 | .3818 |
| PANSS x Timepoint | -0.0003 | [-0.0007, 0.0002] | .2106 | .3285 |
| | | Cingulum Bundle | | |
| PANSS | 0.0003 | [-0.0001, 0.0007] | .1715 | .1715 |
| Timepoint | 0.0027 | [-0.0036, 0.0091] | .4025 | .4025 |
| PANSS x Timepoint | -0.0003 | [-0.0008, 0.0003] | .3285 | .3285 |
| | | Superior Longitudinal Fasciculus | | |
| PANSS | 0.0004 | [-0.0000, 0.0007] | .0529 | .1303 |
| Timepoint | 0.0033 | [-0.0027, 0.0093] | .2863 | .3818 |
| PANSS x Timepoint | -0.0003 | [-0.0008, 0.0002] | .3009 | .3285 |
| | | Inferior Fronto-Occipital Fasciculus | | |
| PANSS | 0.0003 | [-0.0001, 0.0007] | .0953 | .1303 |
| Timepoint | 0.0042 | [-0.0017, 0.0101] | .1643 | .3818 |
| PANSS x Timepoint | -0.0003 | [-0.0008, 0.0001] | .1601 | .3285 |

*Note.* Fixed effects from the secondary analysis linear mixed-effects model (random intercept for subject); testing for effects of PANSS and time on fractional anisotropy (FA), while controlling for age, gender, ethnicity, baseline antipsychotic dose, and baseline IQ. Benjamin-Hochberg correction was applied to adjust for multiple comparisons. None of the outcomes of interest (PANSS, timepoint, PANSS x Timepoint) reached statistical significance.

**Table S9**

*Secondary Analysis Fixed Effects – CAARMS*

| Fixed Effect | est | 95 % CI | *p* | *p* corr |
|---|---|---|---|---|
| | | Global FA | | |
| CAARMS | 0.0001 | [-0.0003, 0.0006] | .5630 | .7571 |
| Timepoint | | | | |
| Month 6 | -0.0009 | [-0.0068, 0.0051] | .7713 | .9268 |
| Month 12 | 0.0021 | [-0.0037, 0.0080] | .4774 | .5389 |
| CAARMS x Timepoint | | | | |
| CAARMS – Month 6 | -0.0000 | [-0.0006, 0.0006] | .9709 | .9709 |
| CAARMS – Month 12 | -0.0002 | [-0.0007, 0.0004] | .5200 | .5328 |
| | | Cingulum Bundle | | |
| CAARMS | 0.0002 | [-0.0004, 0.0008] | .4737 | .7571 |
| Timepoint | | | | |
| Month 6 | -0.0012 | [-0.0087, 0.0062] | .7468 | .9268 |
| Month 12 | 0.0029 | [-0.0044, 0.0102] | .4390 | .5389 |
| CAARMS x Timepoint | | | | |
| CAARMS – Month 6 | -0.0000 | [-0.0007, 0.0007] | .9572 | .9709 |
| CAARMS – Month 12 | -0.0003 | [-0.0010, 0.0004] | .4467 | .5328 |
| | | Superior Longitudinal Fasciculus | | |
| CAARMS | 0.0001 | [-0.0004, 0.0006] | .7571 | .7571 |
| Timepoint | | | | |
| Month 6 | -0.0016 | [-0.0082, 0.0050] | .6306 | .9268 |
| Month 12 | 0.0020 | [-0.0044, 0.0085] | .5389 | .5389 |
| CAARMS x Timepoint | | | | |
| CAARMS – Month 6 | 0.0000 | [-0.0006, 0.0007] | .8890 | .9709 |
| CAARMS – Month 12 | -0.0002 | [-0.0008, 0.0004] | .5260 | .5328 |
| | | Inferior Fronto-Occipital Fasciculus | | |
| CAARMS | 0.0001 | [-0.0004, 0.0006] | .5818 | .7571 |
| Timepoint | | | | |
| Month 6 | -0.0003 | [-0.0068, 0.0062] | .9268 | .9268 |
| Month 12 | 0.0025 | [-0.0039, 0.0088] | .4466 | .5389 |
| CAARMS x Timepoint | | | | |
| CAARMS – Month 6 | -0.0001 | [-0.0007, 0.0005] | .7603 | .9709 |
| CAARMS – Month 12 | -0.0002 | [-0.0008, 0.0004] | .5328 | .5328 |

*Note.* Fixed effects from the secondary analysis linear mixed-effects model (random intercept for subject); testing for effects of CAARMS and time on fractional anisotropy (FA), while controlling for age, gender, ethnicity, baseline antipsychotic dose, and baseline IQ. Benjamin-Hochberg correction was applied to adjust for multiple comparisons. None of the outcomes of interest (CAARMS, timepoint, CAARMS x Timepoint) reached statistical significance.

**Table S10**

*Sensitivity Analysis Fixed Effects – No CPZ*

| Fixed Effect | est | 95 % CI | *p* | *p* corr |
|---|---|---|---|---|
| | | Global FA | | |
| Group | | | | |
| CHR vs HC | -0.0024 | [-0.0073, 0.0024] | .3209 | .7086 |
| FEP vs HC | -0.0060 | [-0.0114, -0.0006] | *.0305 | .0722 |
| Timepoint | | | | |
| Month 6 | 0.0003 | [-0.0019, 0.0025] | .7959 | .9036 |
| Month 12 | -0.0014 | [-0.0036, 0.0008] | .1993 | .3092 |
| Group x Timepoint | | | | |
| CHR – Month 6 | -0.0018 | [-0.0046, 0.0010] | .2044 | .2834 |
| FEP – Month 6 | 0.0016 | [-0.0112, 0.0144] | .8060 | .8060 |
| CHR – Month 12 | 0.0013 | [-0.0016, 0.0041] | .3846 | .5238 |
| FEP – Month 12 | 0.0014 | [-0.0013, 0.0042] | .2940 | .6180 |
| | | Cingulum Bundle | | |
| Group | | | | |
| CHR vs HC | -0.0022 | [-0.0085, 0.0040] | .4794 | .7086 |
| FEP vs HC | -0.0068 | [-0.0138, 0.0002] | .0570 | .0760 |
| Timepoint | | | | |
| Month 6 | 0.0002 | [-0.0025, 0.0029] | .9036 | .9036 |
| Month 12 | -0.0014 | [-0.0041, 0.0013] | .2996 | .3092 |
| Group x Timepoint | | | | |
| CHR – Month 6 | -0.0022 | [-0.0056, 0.0013] | .2126 | .2834 |
| FEP – Month 6 | 0.0062 | [-0.0095, 0.0219] | .4404 | .8060 |
| CHR – Month 12 | 0.0011 | [-0.0023, 0.0046] | .5238 | .5238 |
| FEP – Month 12 | 0.0008 | [-0.0025, 0.0042] | .6180 | .6180 |
| | | Superior Longitudinal Fasciculus | | |
| Group | | | | |
| CHR vs HC | -0.0013 | [-0.0066, 0.0039] | .6269 | .7086 |
| FEP vs HC | -0.0037 | [-0.0096, 0.0022] | .2150 | .2150 |
| Timepoint | | | | |
| Month 6 | 0.0003 | [-0.0021, 0.0028] | .7882 | .9036 |
| Month 12 | -0.0013 | [-0.0037, 0.0012] | .3092 | .3092 |
| Group x Timepoint | | | | |
| CHR – Month 6 | -0.0021 | [-0.0052, 0.0011] | .1994 | .2834 |
| FEP – Month 6 | 0.0079 | [-0.0064, 0.0222] | .2792 | .8060 |
| CHR – Month 12 | 0.0011 | [-0.0021, 0.0042] | .5070 | .5238 |
| FEP – Month 12 | 0.0011 | [-0.0019, 0.0041] | .4720 | .6180 |
| | | Inferior Fronto-Occipital Fasciculus | | |
| Group | | | | |
| CHR vs HC | -0.0010 | [-0.0063, 0.0043] | .7086 | .7086 |
| FEP vs HC | -0.0064 | [-0.0124, -0.0004] | *.0361 | .0722 |
| Timepoint | | | | |
| Month 6 | -0.0005 | [-0.0029, 0.0019] | .6858 | .9036 |
| Month 12 | -0.0015 | [-0.0039, 0.0009] | .2222 | .3092 |
| Group x Timepoint | | | | |
| CHR – Month 6 | -0.0011 | [-0.0042, 0.0020] | .4763 | .4763 |

| | | | | |
|---|---|---|---|---|
| FEP – Month 6 | 0.0027 | [-0.0114, 0.0168] | .7085 | .8060 |
| CHR – Month 12 | 0.0016 | [-0.0015, 0.0047] | .3226 | .5238 |
| FEP – Month 12 | 0.0015 | [-0.0015, 0.0045] | .3166 | .6180 |

*Note.* Fixed effects from the sensitivity analysis linear mixed-effects model (random intercept for subject); testing for effects of group status and time on fractional anisotropy (FA), while controlling for age, gender, ethnicity, and baseline IQ; not for baseline antipsychotic dose as done in the primary analysis. Benjamin-Hochberg correction was applied to adjust for multiple comparisons. None of the outcomes of interest (Group, Timepoint, Group x Timepoint) reached statistical significance.

* $p < .05$

**Table S11**

*Sensitivity Analysis Fixed Effects – Gender: Male*

| Fixed Effect | est | 95 % CI | *p* | *p* corr |
|---|---|---|---|---|
| | | Global FA | | |
| Group | | | | |
| CHR vs HC | -0.0003 | [-0.0069, 0.0063] | .9176 | .9176 |
| FEP vs HC | -0.0033 | [-0.0112, 0.0045] | .4072 | .7174 |
| Timepoint | | | | |
| Month 6 | -0.0006 | [-0.0037, 0.0026] | .7176 | .7176 |
| Month 12 | -0.0015 | [-0.0045, 0.0016] | .3471 | .4595 |
| Group x Timepoint | | | | |
| CHR – Month 6 | -0.0012 | [-0.0053, 0.0029] | .5615 | .8718 |
| FEP – Month 6 | 0.0025 | [-0.0160, 0.0211] | .7895 | .9415 |
| CHR – Month 12 | 0.0022 | [-0.0018, 0.0062] | .2871 | .3725 |
| FEP – Month 12 | 0.0005 | [-0.0032, 0.0042] | .7854 | .9164 |
| | | Cingulum Bundle | | |
| Group | | | | |
| CHR vs HC | -0.0012 | [-0.0096, 0.0073] | .7892 | .9176 |
| FEP vs HC | -0.0028 | [-0.0129, 0.0073] | .5899 | .7174 |
| Timepoint | | | | |
| Month 6 | -0.0014 | [-0.0052, 0.0024] | .4768 | .7176 |
| Month 12 | -0.0018 | [-0.0055, 0.0019] | .3382 | .4595 |
| Group x Timepoint | | | | |
| CHR – Month 6 | -0.0006 | [-0.0056, 0.0044] | .8054 | .8718 |
| FEP – Month 6 | 0.0106 | [-0.0119, 0.0330] | .3565 | .7131 |
| CHR – Month 12 | 0.0031 | [-0.0017, 0.0079] | .2099 | .3725 |
| FEP – Month 12 | 0.0002 | [-0.0042, 0.0047] | .9164 | .9164 |
| | | Superior Longitudinal Fasciculus | | |
| Group | | | | |
| CHR vs HC | 0.0015 | [-0.0059, 0.0089] | .6937 | .9176 |
| FEP vs HC | -0.0016 | [-0.0104, 0.0072] | .7174 | .7174 |
| Timepoint | | | | |
| Month 6 | -0.0007 | [-0.0043, 0.0028] | .6791 | .7176 |
| Month 12 | -0.0013 | [-0.0048, 0.0021] | .4595 | .4595 |
| Group x Timepoint | | | | |
| CHR – Month 6 | -0.0016 | [-0.0062, 0.0031] | .5057 | .8718 |
| FEP – Month 6 | 0.0135 | [-0.0075, 0.0344] | .2083 | .7131 |
| CHR – Month 12 | 0.0021 | [-0.0025, 0.0066] | .3725 | .3725 |
| FEP – Month 12 | 0.0002 | [-0.0039, 0.0044] | .9130 | .9164 |
| | | Inferior Fronto-Occipital Fasciculus | | |
| Group | | | | |
| CHR vs HC | 0.0020 | [-0.0053, 0.0094] | .5868 | .9176 |
| FEP vs HC | -0.0034 | [-0.0122, 0.0053] | .4448 | .7174 |
| Timepoint | | | | |
| Month 6 | -0.0015 | [-0.0050, 0.0019] | .3835 | .7176 |
| Month 12 | -0.0016 | [-0.0049, 0.0018] | .3610 | .4595 |
| Group x Timepoint | | | | |
| CHR – Month 6 | -0.0004 | [-0.0049, 0.0042] | .8718 | .8718 |

| | | | | |
|---|---|---|---|---|
| FEP – Month 6 | 0.0008 | [-0.0197, 0.0212] | .9415 | .9415 |
| CHR – Month 12 | 0.0025 | [-0.0019, 0.0069] | .2610 | .3725 |
| FEP – Month 12 | 0.0004 | [-0.0036, 0.0045] | .8289 | .9164 |

*Note.* Fixed effects from the sensitivity analysis linear mixed-effects model (random intercept for subject); testing for effects of group status and time on fractional anisotropy (FA), while controlling for age, gender, ethnicity, baseline antipsychotic dose, and baseline IQ; split by gender, here: male. Benjamin-Hochberg correction was applied to adjust for multiple comparisons. None of the outcomes of interest (Group, Timepoint, Group x Timepoint) reached statistical significance.

**Table S12**

*Sensitivity Analysis Fixed Effects – Gender: Female*

| Fixed Effect | est | 95 % CI | *p* | *p* corr |
|---|---|---|---|---|
| | | Global FA | | |
| Group | | | | |
| CHR vs HC | -0.0044 | [-0.0116, 0.0027] | .2267 | .3732 |
| FEP vs HC | -0.0011 | [-0.0109, 0.0087] | .8256 | .9317 |
| Timepoint | | | | |
| Month 6 | 0.0015 | [-0.0014, 0.0043] | .3237 | .4316 |
| Month 12 | -0.0014 | [-0.0044, 0.0017] | .3739 | .6399 |
| Group x Timepoint | | | | |
| CHR – Month 6 | -0.0029 | [-0.0066, 0.0008] | .1259 | .1945 |
| FEP – Month 6 | -0.0001 | [-0.0167, 0.0165] | .9900 | .9923 |
| CHR – Month 12 | -0.0001 | [-0.0040, 0.0038] | .9676 | .9676 |
| FEP – Month 12 | 0.0035 | [-0.0005, 0.0074] | .0871 | .1862 |
| | | Cingulum Bundle | | |
| Group | | | | |
| CHR vs HC | -0.0018 | [-0.0112, 0.0075] | .6998 | .6998 |
| FEP vs HC | 0.0006 | [-0.0122, 0.0134] | .9278 | .9317 |
| Timepoint | | | | |
| Month 6 | 0.0023 | [-0.0014, 0.0059] | .2250 | .4316 |
| Month 12 | -0.0008 | [-0.0046, 0.0030] | .6807 | .6807 |
| Group x Timepoint | | | | |
| CHR – Month 6 | -0.0048 | [-0.0094, -0.0001] | *.0469 | .1874 |
| FEP – Month 6 | -0.0001 | [-0.0212, 0.0210] | .9923 | .9923 |
| CHR – Month 12 | -0.0017 | [-0.0066, 0.0032] | .4977 | .9676 |
| FEP – Month 12 | 0.0024 | [-0.0026, 0.0074] | .3422 | .3422 |
| | | Superior Longitudinal Fasciculus | | |
| Group | | | | |
| CHR vs HC | -0.0044 | [-0.0118, 0.0031] | .2537 | .3732 |
| FEP vs HC | 0.0004 | [-0.0097, 0.0106] | .9317 | .9317 |
| Timepoint | | | | |
| Month 6 | 0.0018 | [-0.0014, 0.0049] | .2696 | .4316 |
| Month 12 | -0.0012 | [-0.0045, 0.0021] | .4800 | .6399 |
| Group x Timepoint | | | | |
| CHR – Month 6 | -0.0030 | [-0.0071, 0.0010] | .1459 | .1945 |
| FEP – Month 6 | 0.0002 | [-0.0179, 0.0183] | .9851 | .9923 |
| CHR – Month 12 | -0.0004 | [-0.0047, 0.0038] | .8483 | .9676 |
| FEP – Month 12 | 0.0030 | [-0.0014, 0.0073] | .1822 | .2429 |
| | | Inferior Fronto-Occipital Fasciculus | | |
| Group | | | | |
| CHR vs HC | -0.0043 | [-0.0121, 0.0035] | .2799 | .3732 |
| FEP vs HC | -0.0019 | [-0.0125, 0.0087] | .7251 | .9317 |
| Timepoint | | | | |
| Month 6 | 0.0009 | [-0.0023, 0.0041] | .5877 | .5877 |
| Month 12 | -0.0014 | [-0.0047, 0.0019] | .4106 | .6399 |
| Group x Timepoint | | | | |
| CHR – Month 6 | -0.0024 | [-0.0065, 0.0017] | .2494 | .2494 |

| | | | | |
|---|---|---|---|---|
| FEP – Month 6 | 0.0043 | [-0.0140, 0.0226] | .6447 | .9923 |
| CHR – Month 12 | 0.0003 | [-0.0040, 0.0046] | .9027 | .9676 |
| FEP – Month 12 | 0.0038 | [-0.0006, 0.0081] | .0931 | .1862 |

*Note.* Fixed effects from the sensitivity analysis linear mixed-effects model (random intercept for subject); testing for effects of group status and time on fractional anisotropy (FA), while controlling for age, gender, ethnicity, baseline antipsychotic dose, and baseline IQ; split by gender, here: female. Benjamin-Hochberg correction was applied to adjust for multiple comparisons. None of the outcomes of interest (Group, Timepoint, Group x Timepoint) reached statistical significance.

* $p < .05$

**Table S13**

*Sensitivity Analysis Fixed Effects – Left Hemisphere*

| Fixed Effect | est | 95 % CI | *p* | *p* corr |
|---|---|---|---|---|
| | | Cingulum Bundle | | |
| Group | | | | |
| CHR vs HC | -0.0015 | [-0.0079, 0.0049] | .6494 | .6931 |
| FEP vs HC | -0.0006 | [-0.0087, 0.0074] | .8767 | .9827 |
| Timepoint | | | | |
| Month 6 | 0.0004 | [-0.0026, 0.0033] | .7999 | .7999 |
| Month 12 | -0.0023 | [-0.0052, 0.0007] | .1324 | .2278 |
| Group x Timepoint | | | | |
| CHR – Month 6 | -0.0016 | [-0.0055, 0.0022] | .3961 | .4624 |
| FEP – Month 6 | 0.0013 | [-0.0158, 0.0183] | .8839 | .8839 |
| CHR – Month 12 | 0.0019 | [-0.0019, 0.0057] | .3238 | .4477 |
| FEP – Month 12 | 0.0013 | [-0.0023, 0.0049] | .4914 | .4914 |
| | | Superior Longitudinal Fasciculus | | |
| Group | | | | |
| CHR vs HC | -0.0011 | [-0.0065, 0.0043] | .6931 | .6931 |
| FEP vs HC | -0.0001 | [-0.0068, 0.0066] | .9827 | .9827 |
| Timepoint | | | | |
| Month 6 | 0.0004 | [-0.0021, 0.0029] | .7467 | .7999 |
| Month 12 | -0.0014 | [-0.0040, 0.0011] | .2642 | .2642 |
| Group x Timepoint | | | | |
| CHR – Month 6 | -0.0017 | [-0.0050, 0.0015] | .2959 | .4624 |
| FEP – Month 6 | 0.0124 | [-0.0022, 0.0270] | .0976 | .2929 |
| CHR – Month 12 | 0.0013 | [-0.0020, 0.0045] | .4477 | .4477 |
| FEP – Month 12 | 0.0011 | [-0.0020, 0.0042] | .4708 | .4914 |
| | | Inferior Fronto-Occipital Fasciculus | | |
| Group | | | | |
| CHR vs HC | -0.0011 | [-0.0065, 0.0043] | .6909 | .6931 |
| FEP vs HC | -0.0015 | [-0.0082, 0.0052] | .6676 | .9827 |
| Timepoint | | | | |
| Month 6 | -0.0005 | [-0.0030, 0.0019] | .6664 | .7999 |
| Month 12 | -0.0018 | [-0.0043, 0.0007] | .1519 | .2278 |
| Group x Timepoint | | | | |
| CHR – Month 6 | -0.0012 | [-0.0044, 0.0020] | .4624 | .4624 |
| FEP – Month 6 | -0.0015 | [-0.0159, 0.0130] | .8413 | .8839 |
| CHR – Month 12 | 0.0021 | [-0.0011, 0.0053] | .2042 | .4477 |
| FEP – Month 12 | 0.0014 | [-0.0017, 0.0045] | .3767 | .4914 |

*Note.* Fixed effects from the sensitivity analysis linear mixed-effects model (random intercept for subject); testing for effects of group status and time on fractional anisotropy (FA), while controlling for age, gender, ethnicity, baseline antipsychotic dose, and baseline IQ; split by hemisphere, here: left. Benjamin-Hochberg correction was applied to adjust for multiple comparisons. None of the outcomes of interest (Group, Timepoint, Group x Timepoint) reached statistical significance.

**Table S14**

*Sensitivity Analysis Fixed Effects – Right Hemisphere*

| Fixed Effect | est | 95 % CI | *p* | *p* corr |
|---|---|---|---|---|
| | | Cingulum Bundle | | |
| Group | | | | |
| CHR vs HC | -0.0017 | [-0.0080, 0.0047] | .6013 | .7601 |
| FEP vs HC | -0.0013 | [-0.0093, 0.0066] | .7432 | .8582 |
| Timepoint | | | | |
| Month 6 | -0.0000 | [-0.0028, 0.0027] | .9786 | .9786 |
| Month 12 | -0.0006 | [-0.0034, 0.0022] | .6712 | .6712 |
| Group x Timepoint | | | | |
| CHR – Month 6 | -0.0033 | [-0.0069, 0.0003] | .0764 | .1806 |
| FEP – Month 6 | 0.0112 | [-0.0051, 0.0274] | .1782 | .5345 |
| CHR – Month 12 | 0.0004 | [-0.0032, 0.0041] | .8124 | .8124 |
| FEP – Month 12 | 0.0003 | [-0.0031, 0.0037] | .8686 | .8686 |
| | | Superior Longitudinal Fasciculus | | |
| Group | | | | |
| CHR vs HC | -0.0010 | [-0.0064, 0.0045] | .7237 | .7601 |
| FEP vs HC | -0.0006 | [-0.0074, 0.0062] | .8582 | .8582 |
| Timepoint | | | | |
| Month 6 | 0.0003 | [-0.0023, 0.0029] | .8438 | .9786 |
| Month 12 | -0.0011 | [-0.0037, 0.0015] | .4015 | .6022 |
| Group x Timepoint | | | | |
| CHR – Month 6 | -0.0027 | [-0.0061, 0.0007] | .1204 | .1806 |
| FEP – Month 6 | 0.0033 | [-0.0119, 0.0185] | .6693 | .6693 |
| CHR – Month 12 | 0.0009 | [-0.0025, 0.0043] | .5881 | .8124 |
| FEP – Month 12 | 0.0009 | [-0.0023, 0.0041] | .5941 | .8686 |
| | | Inferior Fronto-Occipital Fasciculus | | |
| Group | | | | |
| CHR vs HC | -0.0009 | [-0.0064, 0.0047] | .7601 | .7601 |
| FEP vs HC | -0.0040 | [-0.0109, 0.0029] | .2598 | .7795 |
| Timepoint | | | | |
| Month 6 | -0.0005 | [-0.0030, 0.0021] | .7244 | .9786 |
| Month 12 | -0.0012 | [-0.0038, 0.0013] | .3455 | .6022 |
| Group x Timepoint | | | | |
| CHR – Month 6 | -0.0014 | [-0.0047, 0.0019] | .4196 | .4196 |
| FEP – Month 6 | 0.0069 | [-0.0079, 0.0217] | .3617 | .5426 |
| CHR – Month 12 | 0.0012 | [-0.0022, 0.0045] | .4936 | .8124 |
| FEP – Month 12 | 0.0015 | [-0.0017, 0.0046] | .3565 | .8686 |

*Note.* Fixed effects from the sensitivity analysis linear mixed-effects model (random intercept for subject); testing for effects of group status and time on fractional anisotropy (FA), while controlling for age, gender, ethnicity, baseline antipsychotic dose, and baseline IQ; split by hemisphere, here: right. Benjamin-Hochberg correction was applied to adjust for multiple comparisons. None of the outcomes of interest (Group, Timepoint, Group x Timepoint) reached statistical significance.

**Table S15**

*Medication and Clinical Scores at Baseline Stratified by Gender in FEP*

| | **Male** (*n* = 108) | **Female** (*n* = 44) |
|---|---|---|
| CPZ (mg/day) | 254 (208) | 155 (166) |
| SOFAS | 53.5 (18.8) | 60.4 (21.2) |
| PANSS – Positive | 13.1 (5.8) | 11.5 (5.8) |

*Note.* Scores in mean (SD).

**Figure S1**

*Correlation of SOFAS Scores and Antipsychotic Dose at Baseline*

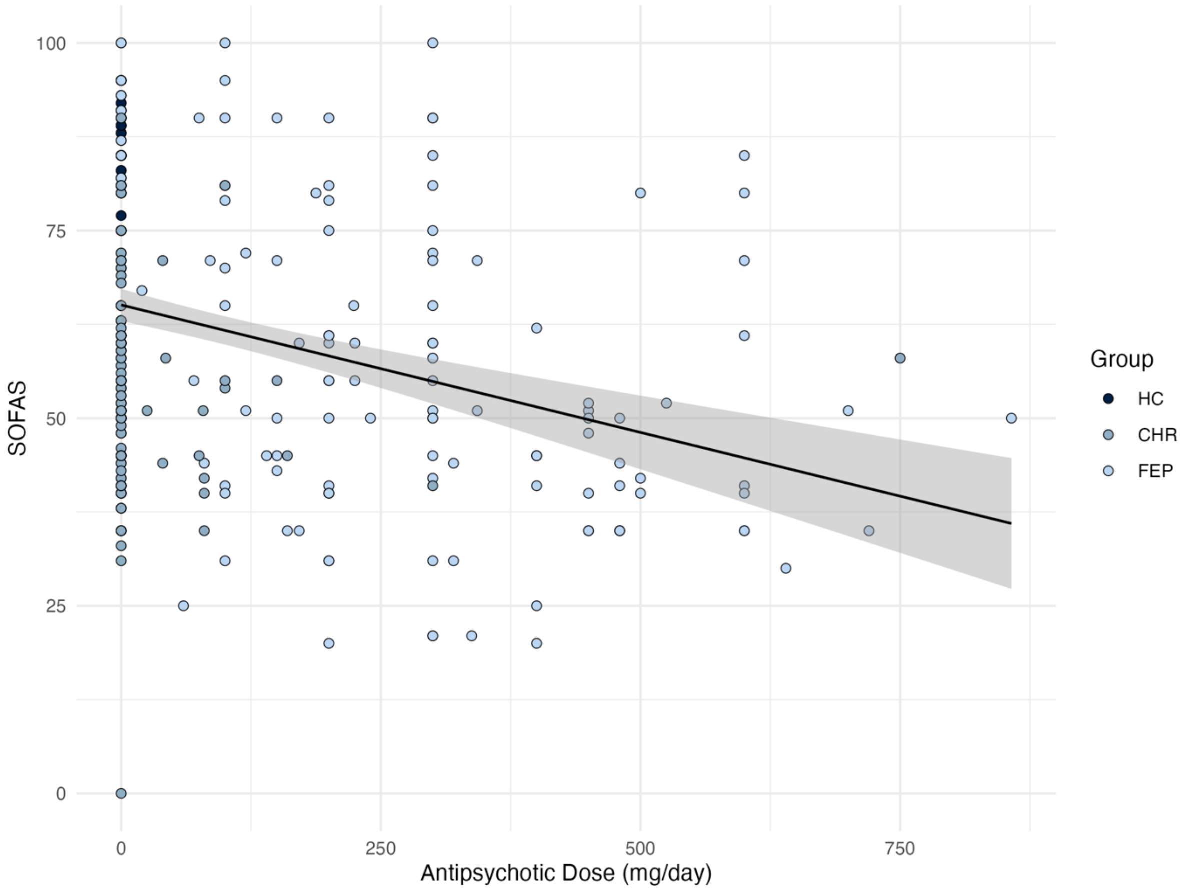

**Figure S2**

*Correlation of PANSS Positive Scores and Antipsychotic Dose at Baseline in FEP*

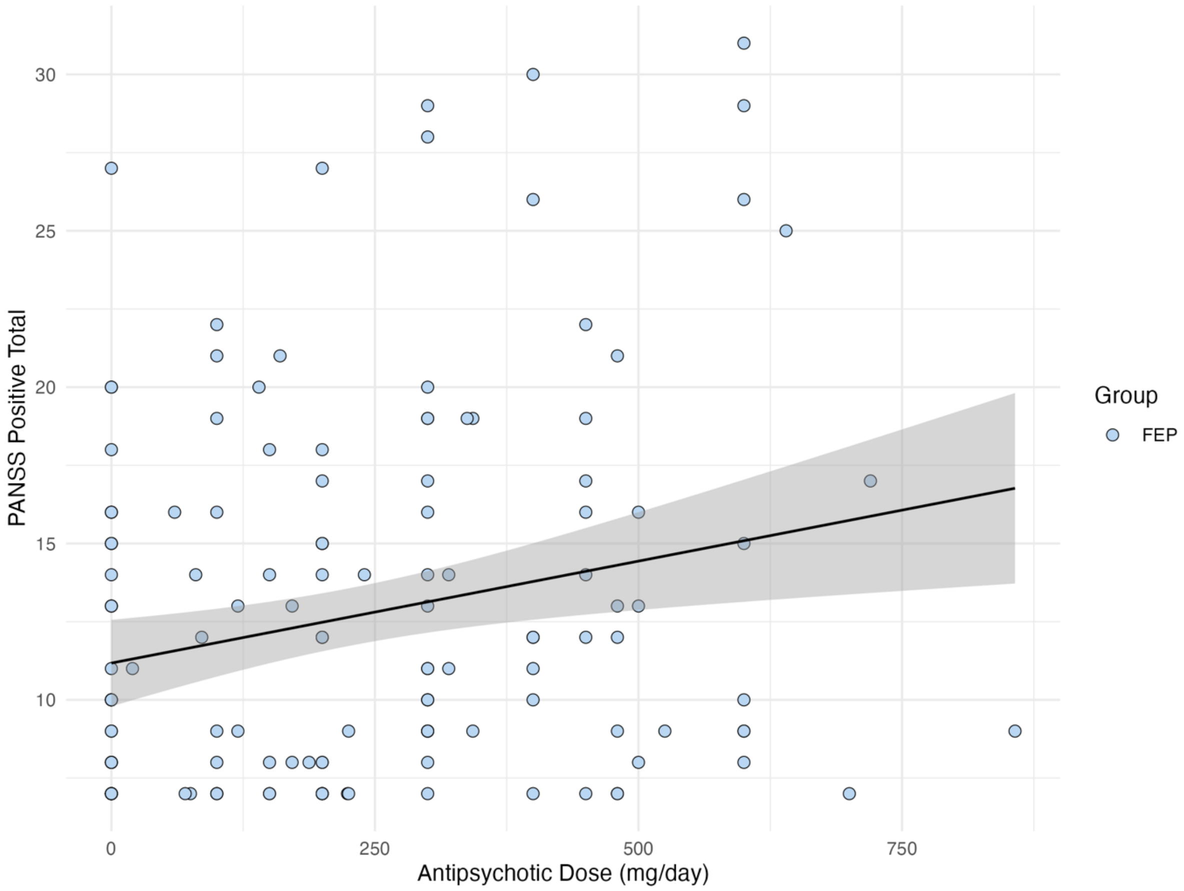
